# Cerebellar contributions to a brainwide network for reversal learning

**DOI:** 10.1101/2021.12.07.471685

**Authors:** Jessica L. Verpeut, Silke Bergeler, Mikhail Kislin, F. William Townes, Ugne Klibaite, Zahra M. Dhanerawala, Austin Hoag, Caroline Jung, Junuk Lee, Thomas J. Pisano, Kelly M. Seagraves, Joshua W. Shaevitz, Samuel S.-H. Wang

**Author notes:** Corresponding authors Correspondence: Samuel S.-H. Wang Neuroscience Institute, Washington Road Princeton University Princeton, New Jersey 08544 USA Jessica Verpeut Department of Psychology Arizona State University 950 South McAllister Avenue Tempe, Arizona 85281 USA.

## Abstract

The cerebellum regulates nonmotor behavior, but the routes by which it exerts its influence are not well characterized. Here we report a necessary role for the posterior cerebellum in guiding a reversal learning task, acting through a network of diencephalic and neocortical structures. After chemogenetic inhibition of Purkinje cells in lobule VI or crus I, mice could learn a water Y-maze task but were impaired in their ability to reverse their initial choice. To map targets of perturbation, we imaged c-Fos activation in cleared whole brains using light-sheet microscopy. Reversal learning activated diencephalic regions and associative neocortical regions. Distinctive subsets of structures were altered by perturbation of lobule VI (including thalamus and habenula) and crus I (including hypothalamus and prelimbic/orbital cortex), and both perturbations influenced anterior cingulate and infralimbic cortex. To identify functional networks, we used correlated variation in c-Fos activation within each test group. Lobule VI inactivation weakened within-thalamus correlations, while crus I inactivation divided neocortical activity into well-separated sensorimotor and associative subnetworks. In both groups, high-throughput automated analysis of complex whole-body movement revealed deficiencies in across-day adaptation to an open field environment. Neither perturbation affected gait, within-day open-field adaptation, or location preference. Taken together, these experiments reveal brainwide systems for cerebellar influence that can affect multiple flexible responses.

**Significance statement:** The cerebellum, a part of all vertebrate brains, provides feedback to the rest of the brain on a split-second basis in response to unexpected events. The consequent changes can range from minute adjustments in movement to broad changes in behavior. In mice, we investigated cerebellar regions that help guide flexible behavior. Inactivation of these regions prevented mice from responding flexibly to a changing Y-shaped maze. We used light-sheet microscopy of c-fos protein expression to map brainwide effects of different stages of learning and cerebellar inactivation. Inactivation caused changes in thalamocortical activity that were similar in pattern, but opposite in sign, from normal learning. Lobule VI inactivation weakened thalamic functional networks, while crus I inactivation divided sensorimotor from associative neocortical networks. The same inactivation also impaired multiday adaptation to an open arena, as tracked using machine vision. Our work shows how the cerebellum’s influence over flexible response can be mediated by a widespread brain network.

## Introduction

The cerebellum is increasingly appreciated for its contributions to flexible behavior, in addition to its better-known role in shaping movement and balance. Prominent anatomical pathways between cerebellum and neocortex suggest a role in higher-order processing [1–5]. In humans, insult to the posterior cerebellum results in a clinical cognitive-affective syndrome that includes impairments in executive function, working memory, abstract reasoning, and emotional processing [6,7]. More severe outcomes arise from pediatric cerebellar insult, including a diagnosis of autism, a disorder characterized by inflexibility to the point of emotional distress when routines are violated [8–3]. Taken together, these studies suggest that, like the neocortex, the cerebellum plays a necessary role in flexible behavior and cognitive processing.

Animal experiments have identified specific regions of the cerebellar cortex that support flexible behavior. In lobule VI, a midline posterior structure that is perturbed in autism spectrum disorder [14,5], inhibition of molecular layer interneurons alters reversal learning, perseverative or repetitive behavior, novelty-seeking, and social preference [16]. Perturbation of rodent crus I, the human homolog [17] of which is structurally altered in ASD, leads to deficits in social, repetitive, and flexible behaviors [16,8], and neither perturbation affects gait. Furthermore, inactivation of Purkinje cells in rodent crus I reduces the ability to perform sensory evidence accumulation, a task in which Purkinje cells have been found to encode choices and accumulated evidence [19,20].

Lobule VI and crus I engage with the forebrain through bidirectional polysynaptic pathways [21]. Purkinje cells in the cerebellar cortex receive input from distal forebrain structures and transsynaptic tracing in mice has identified inhibitory output to cerebellar and vestibular nuclei, which in turn provide excitatory output to the rest of the brain forming the cerebral-thalamic-cerebellar circuit [1–4,22,23]. Along these pathways, cerebellar cortex is organized into parasagittal microzones which project in distinctive patterns, so that lobule VI and crus I make different patterns of disynaptic connectivity with thalamic structures [24–6], and trisynaptic paths to anterior cingulate, infralimbic, and somatosensory cortex [1–5]. Each of these cerebellar regions also receives descending input from the neocortex via the pons [27,8] and inferior olive [29]. These cerebellar regions therefore have distinctive routes by which they may influence forebrain processing across many distributed targets.

To interrogate the contribution of the posterior cerebellum to a simple reversal learning task, we monitored mouse behavior and mapped brain-wide patterns of activation after perturbing lobule VI and crus I. First, we chemogenetically perturbed neural activity reversibly in Purkinje cells, the principal output neurons of the cerebellar cortex. Second, we combined a Y-maze learning paradigm with c-Fos mapping to identify brain-wide substrates of reversal learning. Third, we studied expression of the activity-dependent gene product c-Fos using tissue clearing techniques combined with light-sheet microscopy to map across the whole brain without need for tissue sectioning. We analyzed this data to identify individual activated regions and patterns of coactivation. Lastly, we characterized freely-moving mouse behavior in granular detail using machine learning methods for automated tracking of body poses, movements, and actions. Together, these approaches provide a framework for characterizing how the cerebellum influences brainwide functional networks to modulate flexible behavior.

## Results

### Experimental design to reversibly perturb Purkinje cells

To probe the impact of cerebellar activity on flexible behavior and whole-brain activity, we chemogenetically inhibited Purkinje cells. Purkinje cells influence the rest of the brain via their projections to the deep cerebellar nuclei (DCN), which send excitatory output to the rest of the brain. The inhibitory DREADD (Designer Receptor Exclusively Activated by Designer Drugs) hM4Di was expressed in Purkinje cells using an adeno-associated virus (AAV) containing the hM4Di sequence under control of the L7 promoter. DREADD expression was robust and confined to Purkinje cells (Fig 1A **and S1 Fig**). In slices, application of the DREADD agonist clozapine-N-oxide (CNO; 10 µM) reduced evoked action potential firing in hM4Di-positive Purkinje cells in slices (paired t-test, p = 0.0009) (Fig 1B-D), thus removing modulation of DCN neurons. To test the consequences for deep-nuclear firing, we made many-electrode recordings in 5 awake mice and identified a total of 43 units in which tactile airpuff stimuli evoked at least a 5 Hz facilitation increase in firing (Fig 1E-G **and S2 Fig**). In these units, application of CNO increased spontaneous firing rates (-10 minutes relative to CNO: 32.5 Hz, SD: 17.6; 20: 36.8 Hz, SD: 18.9; 50: 39.0 Hz, SD: 22.5; paired t-test: p = 0.033 -10 vs 20 min, p = 0.013 20 vs 50 min) (Fig 1F **and S2 Fig**). Before CNO application, tactile airpuffs first evoked an increase in DCN activity during the airpuff, consistent with mossy-fiber and/or climbing-fiber drive, followed by a strong undershoot. Inhibition of Purkinje cells abolished the undershoot, leaving a sustained increase in firing that outlasted the duration of the airpuff, consistent with decreased spontaneous Purkinje cell drive and consequent disinhibition of DCN neurons (-10 min: 32.5 +/-17.6, 20: 36.8 +/-18.9, 50: 39 +/-22.5 Hz).

**Fig 1.**
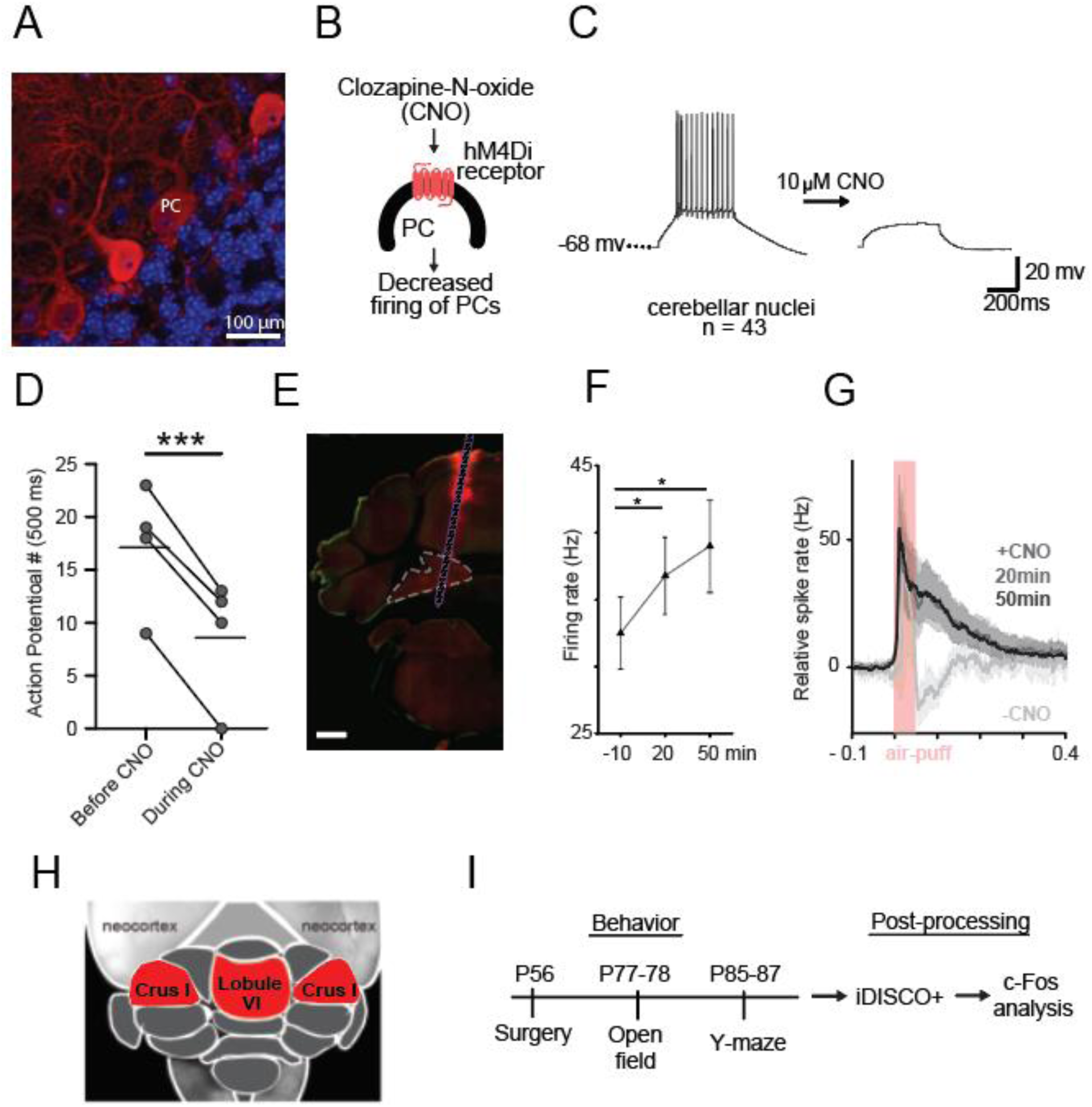
Acute adult inactivation of Purkinje cells in the cerebellar cortex. (A) Expression of the chemogenetic DREADD probe in Purkinje cells (red). (B) The activating ligand CNO binds to the hM4Di receptor and decreases Purkinje cells firing. (C) Slice electrophysiology of Purkinje cells before and during CNO (10 μM). (D) Purkinje cells with DREADD receptors fire fewer action potentials in response to current injection. (E) Example histological section with the probe location marked by CM-DiI (red), scale bar, 0.5 mm. (F) Average spontaneous firing rate of deep cerebellar nuclei neurons increases after activation of DREADD receptors in Purkinje cells. (G) Average response (averaged across all sensory stimulation and all sorted units, background subtracted and normalized) in deep cerebellar nuclei (mean ± confidence interval) before then 20 and 50 min after CNO injection (n = 5 mice). (H) Dorsal view of the cerebellum with targeted lobule VI or crus I (red). (I) Adult mice (PND 56) received surgery for AAV injection (DREADDs or mCherry) at PND 56 for behavioral testing starting with open field between PND 77-78 and water Y-maze starting between PND 85-87. For each behavior test, animals received CNO (i.p.), vehicle, or no treatment 20 minutes prior to testing. After behavioral testing, brains were cleared for light-sheet microscopy using iDISCO+ and analysis of c-Fos immunopositive cells. Comparisons were made using a paired t-test. * p<0.05, *** p < 0.001

Posterior vermis (lobule VI and VII) and ansiform area (crus I and crus II) have been implicated in non-motor executive functions (Fig 1H) [16,30–32]. We stereotaxically injected virus into either lobule VI or crus I (Allen Brain Atlas ansiform area) at postnatal day (PND) 56 and at sacrifice quantified mCherry-positive voxels (**S1 Fig**). Expression arising from injection of these two targets was restricted to the target, with principal spillover into lobule VII and crus II, respectively (**S1 Fig**). We administered CNO (1 mg/kg intraperitoneal) on test days between PND 77 and PND 90 and tested animals on two paradigms for assessing flexible behavior: reversal learning in a water Y-maze, and spontaneous behavior in an open-field arena (**Table 1**). At different points of Y-maze training, we sacrificed mice and collected tissue for whole-brain imaging of the activity-dependent immediate early gene c-Fos (Fig 1I).

**Table 1.** Experimental groups denoting DREADD and CNO conditions. Animals underwent surgery for an injection of DREADD or mCherry AAV expression. Prior to open field and the water Y-maze task, animals were injected with the DREADD ligand (CNO), Vehicle, or remained untreated. * Denotes previous data from [42].

| Group | DREADDs | CNO | Open Field | Y-maze |
| --- | --- | --- | --- | --- |
| Untreated* | no | no | 60 | 63 |
| Vehicle only | no | no | 9 | 9 |
| CNO only | no | yes | 12 | 22, 7: no reversal |
| mCherry | no | yes | 10 | 18 |
| Lobule VI | yes | yes | 10 | 16, 10: no reversal |
| Bilateral Crus I | yes | yes | 10 | 7 |
| Crus I right | yes | yes | 10 | 25 |
| Crus I left | yes | yes | 10 | 26 |
| L7Cre; Tsc1<sup>flox/flox</sup>* | no | no | 9 | 0 |

### Cerebellar disruption of lobule VI or bilateral crus I impairs Y-maze reversal learning

To test flexible learning, we trained mice in a water Y-maze (Fig 2A). After 1 day of habituation to the environment, an underwater platform was placed at the end of one Y-arm and the mice spent two days learning to find the platform through trial and error (acquisition days 2 and 3). On day 4, the platform was switched to the opposite arm for 4 sessions (reversal). On the fifth and final session on day 4, a barrier was placed blocking the originally learned side (forced reversal). On all days, we defined the correct choice as an entrance to the correct arm and climbing onto the platform during a 40-second trial.

**Fig 2.**
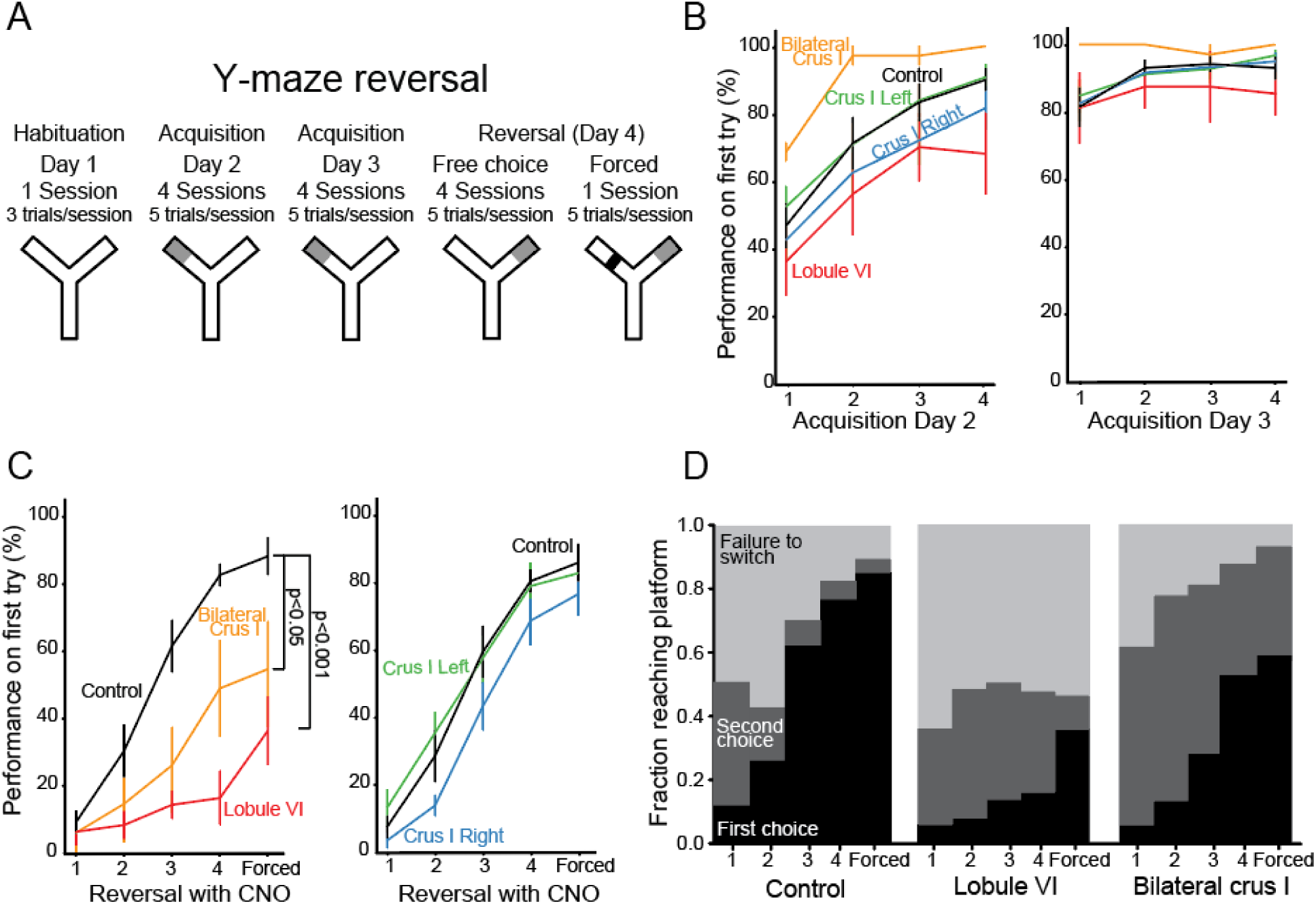
Effect of lobule VI and crus I inactivation on behavior. (A) Protocol for water Y-maze reversal consisting of habituation, two days of training (day 2 and day 3), and one reversal day ending in a forced session (day 4). (B) Animals learned to find the platform over two days. First try performance of 80% at the end of session 4 was required to continue to reversal. (C) Y-maze reversal was impaired in animals with lobule VI and bilateral crus I (left) inactivation compared to CNO only control. Unilateral effects were not found (right) compared to controls. (D) Typical mice learned through trial and error, demonstrating more accurate performance over time. Lobule VI and crus I perturbed mice stayed repeatedly with the original incorrect choice compared to CNO only control. Error bars indicate mean ± SEM. Significance values in (C) indicate comparisons using a generalized mixed effect model (GLMM) with a binomial distribution.

All DREADD-activated groups showed a similar time course of initial acquisition, showing no statistically detectable differences compared with controls (generalized linear mixed-effect model, GLMM, p=0.76; Fig 2B; for control experimental design, see **Methods**). Control treatments did not affect distance swum, initial learning, or reversal learning compared with untreated mice (**S3 Fig**). Lobule VI-perturbed mice and bilateral crus-I-perturbed mice were strongly impaired in reversal learning (Fig 2C **and S3E Fig)**; lobule VI compared to CNO-without-DREADD controls (p = 8.9*10^-6^, Cohen’s *d* = 3.84 and bilateral crus I p = 0.04, Cohen’s *d* = 1.93). Performance on the forced-reversal session was also reduced (lobule VI, p = 0.0039) (**S1 Movie**). In tests for lateralization of crus I function [33], we found that perturbation by crus I injection on either left (p = 0.83) or right (p = 0.10) side alone was insufficient to impair reversal (Fig 3C).

**Fig 3.**
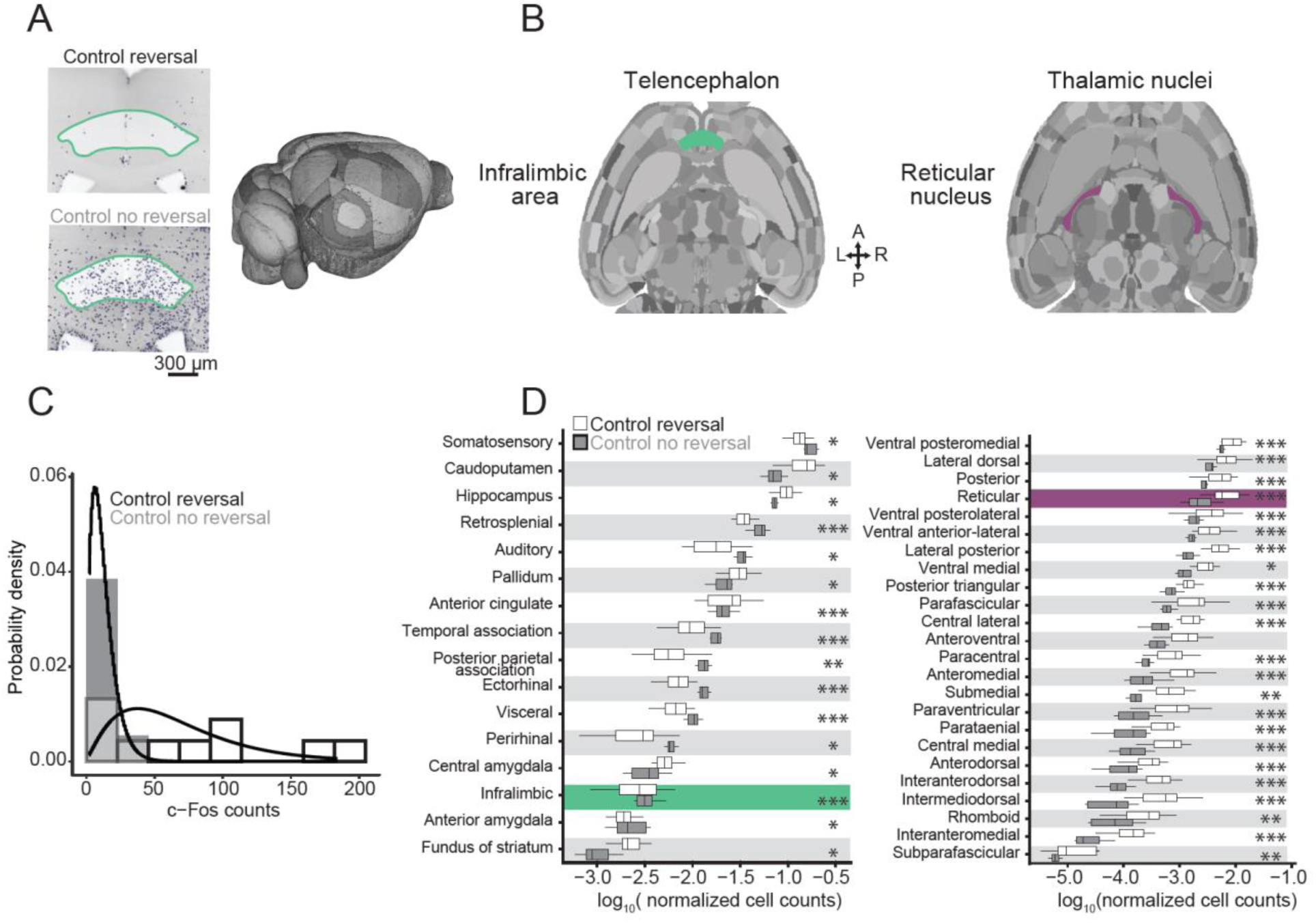
Whole-brain analysis of c-Fos in CNO only controls. (A) Example of ClearMap cell counting in the infralimbic area in CNO only reversal compared to CNO only no reversal (left). Neuroglancer 3-D mouse brain (right). (B) Example horizontal mouse brain sections representing infralimbic area (green) and reticular nucleus (purple). (C) Negative binomial regression under two CNO only controls undergoing reversal or no reversal. This example, nucleus raphe pontis. Nonparametric kernel density estimates derived from count-based densities (bars). Solid curves indicate the probabilities at each count level from a fitted negative binomial regression model. (D) Normalized cell counts (log_10_) in the telencephalon (left) and thalamus (right) comparing CNO only reversal (white) and no reversal (gray). Comparisons were made using a negative binomial regression. * p < 0.05, ** p < 0.01, *** p < 0.001

To probe behavioral patterns of this learning failure, we analyzed individual trials. Entrance into any arm N times, ending with a landing in the correct arm, was defined as an N-th choice trial. Even in the first reversal session, control mice (CNO only) typically found the platform in their first or second choice (85% of mice), eventually making a correct first choice in 71% of trials by the fourth session. In contrast, lobule-VI-perturbed mice made persistent errors even by the fourth session, making correct first choices on 16% of trials and correct second choices on 34% of trials (Fig 2D **and S3F Fig**). In forced-reversal trials, lobule-VI-perturbed mice displayed a unique perseverative behavior of swimming back and forth between the divider and beginning of the maze instead of switching to the obvious open arm, resulting in animals failing to switch in 64% of forced-reversal trials (**S1 Movie**). We also observed perseveration in crus-I perturbed mice, whereby mice continued to swim to the previously correct arm and failed to switch in 47% of forced-reversal trials. (Fig 2D **and S3F Fig**). In summary, mice typically learned the Y-maze by trying multiple arms until the platform was found, but perturbing the cerebellum resulted in persistent errors and a failure to reverse.

### Whole-brain c-Fos reveals lobule-specific targets of influence

We next sought to identify brain-wide targets of cerebellar influence that could account for the observed perseverative and inflexible behavior. To quantify expression of the activity-dependent immediate early gene c-Fos, we extracted brains at different points in the Y-maze learning and reversal paradigm [34]. We compared the effects of lobule VI-targeted and crus I-targeted perturbations with closely-matched control conditions (see **Methods** for description). Brains were cleared using iDISCO+ [35] and immunostained for c-Fos and the mCherry fluorescent tag encoded by both DREADD and control AAVs (**S2 Movie**). Samples were imaged for AlexaFluor-647 on a light-sheet microscope, aligned to the Princeton Mouse Atlas [1–5,36], and analyzed for c-Fos positive cells using ClearMap (Fig 3A **and S2 Movie**) [37]. Using 122 chosen structures, a 3-D representation of the data was created for each brain using Neuroglancer, a Google WebGL-based viewer for volumetric data (Fig 3A) for analysis of c-Fos in single or combined regions (Fig 3B).

Within each paired comparison, we processed brains from all control and treatment animals as a single batch using the same tissue preparation and imaging conditions whenever possible. In the few cases where we needed multiple batches, we adjusted for confounding batch effects by including indicator variables as covariates in regression models. Analogous to genome-wide association studies, we analyzed each brain region independently and corrected the results *post hoc* for false discovery rate. For each region, we quantified the contrast between counts for the animals in the treatment group versus the control group using negative binomial regression with a log link function. To account for animal-specific variation in total count, we included the log of total counts as an offset (Fig 3C).

To identify brain regions activated in the initial acquisition of Y-maze learning, we assayed c-Fos-positive cells immediately after 3 days of Y-maze ending on the last initial acquisition day (n=10 mice), using for baseline comparison animals that underwent habituation-only on the first day of Y-maze (n=10 mice). Out of 122 regions, 33 regions showed increased activity and 6 regions showed decreased activity. We found activation in thalamus (1.4-fold) and in prelimbic (1.8-fold) and temporal association (1.65-fold) cortex. In the rest of the brain, we found some of the strongest associations in parabrachial nucleus (2.8-fold), basolateral, central, and cortical amygdalar nucleus (1.8, 1.7, and 1.7-fold), lateral habenula (2.3-fold), periaqueductal gray (2.1- fold), septohippocampal nucleus (2.9-fold), and lateral septal nucleus (2.6-fold) (**S5: left Fig**). These increases occurred despite a 57% decrease in distance swum (first day, 54.6±4.4 cm; third day, 23.4±6.2 cm, mean±SD) (**S1 Table)**. We confirmed results in several of these regions using conventional immunohistochemistry (p = 0.00006, Cohen’s *d* = 1.4) (**S7A-C Fig, S8 Fig)**. Overall, initial learning specifically activated a wide range of regions linked with associative and affective function.

We analyzed specific neural correlates of reversal learning by comparing brains on day 4 after one day of reversal learning (n=10 mice) with mice undergoing a third day of acquisition (n=8). The distance swum between these conditions differed by a factor of 1.2 (**S1 Table**). Reversal learning resulted in 16 decreased and 44 increased structures compared to the third day of acquisition. We found statistically significant activation throughout thalamus (2.7-fold), including polymodal regions (3.2-fold) as classified by Jones [38], sensory/motor regions (2.1- fold), and the reticular nucleus, which is modulatory (3.2-fold) [38,9]. Reversal learning was associated with decreases in c-Fos cell counts in the majority of neocortical regions, the largest change being a decrease in infralimbic cortex activity (0.56-fold). Additional activation occurred in medial and lateral habenula (15.6-fold and 7.6-fold), periaqueductal gray (1.9-fold), and parabrachial nucleus (1.6-fold) (Fig 3, Fig 5A: **middle, S4 Fig, S6: middle Fig, and S8 Fig**).

We tested the effects of lobule VI-targeted perturbation (n=10 mice) on c-Fos generation in reversal learning. Perturbation of lobule VI during reversal learning resulted in a reduction in c-Fos activity throughout the thalamus (overall 0.28-fold compared with unperturbed reversal learning). The distance swum between lobule-VI perturbation and unperturbed reversal differed by a factor of 1.1 (**S1 Table**). Increased activity was seen in somatomotor, somatosensory, anterior cingulate, and infralimbic cortex. Midbrain regions both increased (ventral tegmental area, lateral hypothalamus, midbrain raphe nuclei) and decreased (parastrial, medial and lateral habenula, and periaqueductal gray) in activity (Fig 4A-C, Fig 5A**: right and C, S6: right Fig, S7D-F Fig, and S8 Fig)**. Thalamic results were confirmed using conventional immunohistochemistry (p = 0.000002, Cohen’s *d* = 2.3) (**S7D-E Fig)**. Such widespread changes were not seen when giving CNO during an additional day of acquisition (Fig 4D).

**Fig 4.**
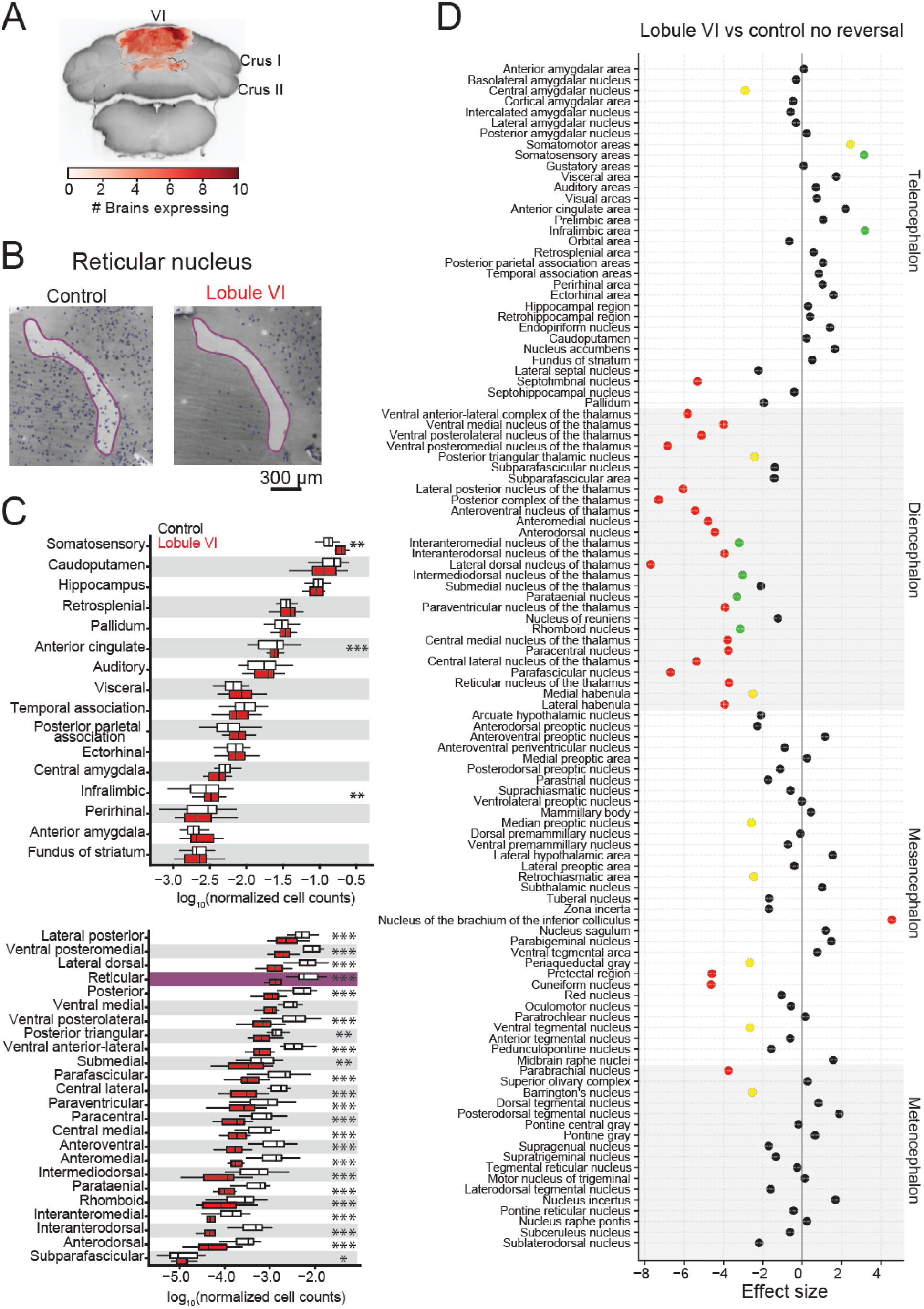
Effects of lobule VI perturbation on whole-brain c-Fos. (A) Overlay of regions of DREADD-AAV expression recovered using immunofluorescent labeling of mCherry and lightsheet microscopy imaging. (B) Example images of ClearMap cell counting in the reticular nucleus comparing Lobule VI and CNO only control after reversal. (C) Normalized cell counts (log_10_) by total regions analyzed in the telencephalon (top) and thalamus (bottom) comparing Lobule VI and CNO only after reversal. (D) Statistically-significant (p < 0.05) lobule VI reversal versus lobule VI control no reversal. Lobule VI perturbation modulated regions necessary for reversal learning, including thalamus. Comparisons were made using a negative binomial regression. Yellow: * p < 0.05, Green: ** p < 0.01, Red: *** p < 0.001

We also examined the consequences of bilateral crus I-targeted perturbation (n = 7) on reversal learning. The distance swum between bilateral crus I-targeted perturbation and reversal learning differed by a factor of 1.3, with bilateral crus I-targeted swimming more (**S1 Table**). Bilateral perturbation of crus I during reversal learning did not lead to statistically detectable differences in thalamic activity compared with unperturbed reversal learning. However, differences were seen in the neocortex, both increased (anterior cingulate, prelimbic, infralimbic, and orbital) and decreased (auditory, visual, posterior parietal, and temporal) activity. Increases were also seen in parastrial nucleus and hypothalamus (lateral and preoptic) (Fig 5A**: left and B and S6: left Fig**). These changes were not seen after unilateral perturbation of crus I (crus I right n = 25, crus I left n = 26) (**S5: middle and right Fig**).

**Fig 5.**
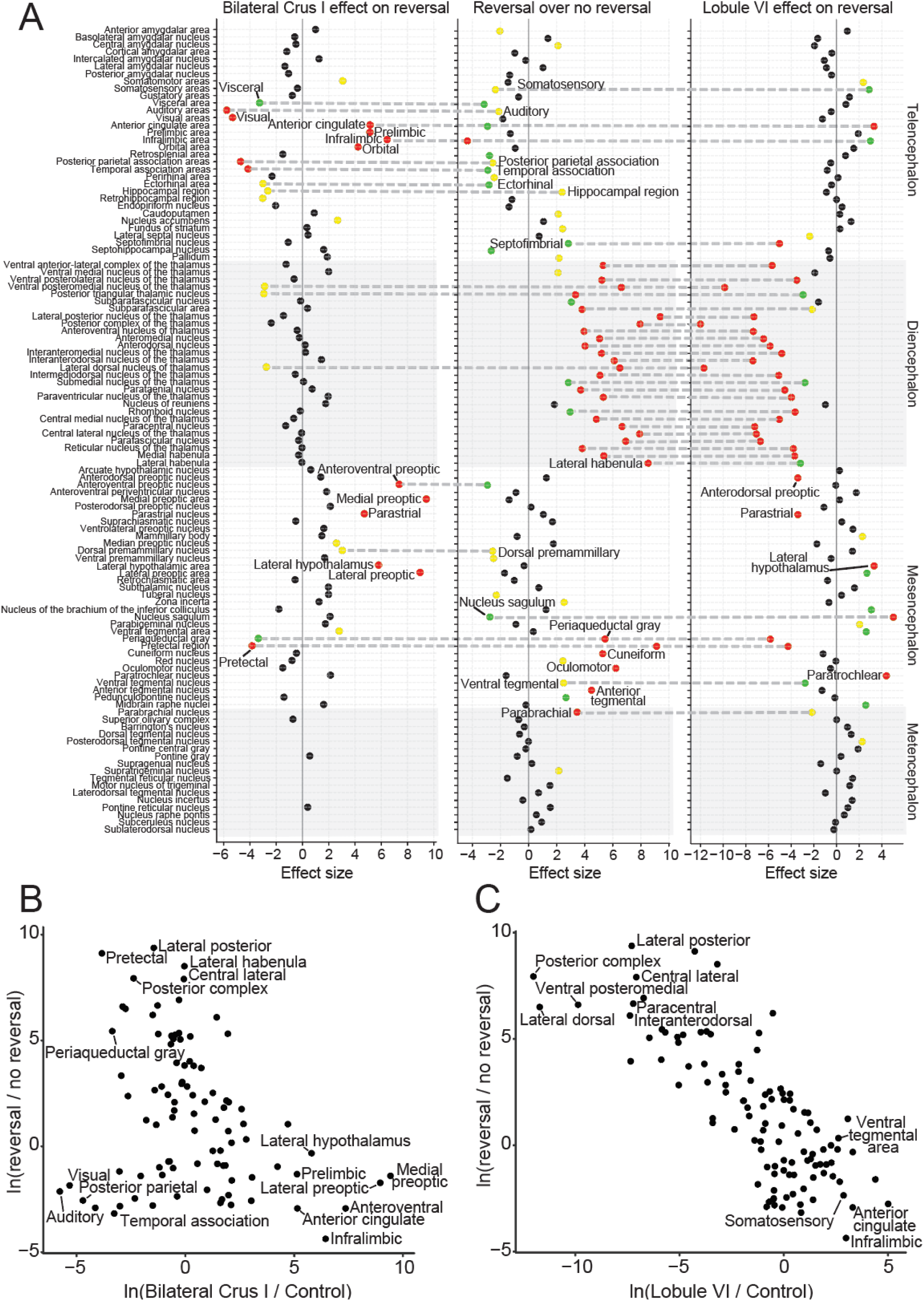
Brain-wide association study (BWAS) to identify activated regions of c-Fos expression. (A) Statistically-significant (p < 0.05) bilateral crus I (left), reversal (middle) lobule VI (right) structures compared to CNO only controls. Lobule VI (right) perturbation modulated regions necessary for reversal learning, including thalamus and anterior cingulate cortex. Crus I bilateral (left) perturbation altered regions in the telencephalon, including prelimbic, infralimbic, anterior cingulate, and orbital frontal cortex. (B) Correlations between crus I and reversal learning structures (Pearson’s r = -0.29). (C) A strong negative correlation was found between lobule VI and reversal learning structures (Pearson’s r = -0.78). All data were calculated as the natural log of ratios rescaled by the standard error. Comparisons were made using a negative binomial regression. Bilateral Crus I and lobule VI were compared to CNO only controls. Yellow: p < 0.05, Green: p < 0.01, Red: p < 0.001

Many effects of lobule VI and crus I perturbation went in the opposite direction as the reversal-versus-acquisition condition. Of the 61 regions showing changed activity in reversal learning, opposite-direction changes occurred in 47 regions from lobule VI perturbation, and in 27 regions from crus I bilateral perturbation. Opposite-direction changes encompassed the majority of thalamic regions and lateral and medial habenula, as well as selected regions in telencephalon (anterior cingulate, infralimbic) and mesencephalon (periaqueductal gray, pretectal regions) with a lobule VI perturbation (Fig 5A). The overall pattern of cell ratios was strongly correlated with the reversal-versus-acquisition condition for lobule VI (correlation of log-cell-ratios by Pearson’s r = -0.78) (Fig 5C), and less correlated for bilateral crus I (Pearson’s r =-0.29) (Fig 5B). Thus, lobule VI perturbation reversed activation patterns in most regions that were activated by reversal learning, while crus I perturbation had effects that were limited to neocortical and hypothalamic regions.

### Reversal learning activates functional networks that are disrupted by cerebellar perturbation

To examine functional connectivity using our c-Fos data, we calculated the Spearman’s rank coefficient ρ among the mice within any one experimental group [40,1]. For crus I, to attain a large sample size we combined left, right, and bilateral inhibition groups. Compared with the reversal learning group, lobule VI and crus I perturbation conditions had stronger within-group correlations in mesencephalon and metencephalon, and weaker correlations within diencephalon and telencephalon (**S9-11 Fig**). This pattern suggested that under either condition, cerebellar perturbation disrupted the coupling between thalamocortical processing and the rest of the brain. To further identify patterns of change, we restricted analysis to structures found to be significant in brainwide comparisons of groups at p < 0.05 in Fig 5: for reversal learning, compared with initial acquisition, and for lobule VI and crus I groups, compared with unperturbed reversal learning (Fig 6 **and S12 Fig**). To visualize the overall pattern of correlation, we used multidimensional scaling to reduce the matrix of correlations to points in a plane, where each point represents one brain region. The more correlated the activity between two regions, the closer the points appear in the plot (Fig 6**: right**). Points were joined by line segments if the absolute value of the correlation was greater than 0.5.

**Fig 6.**
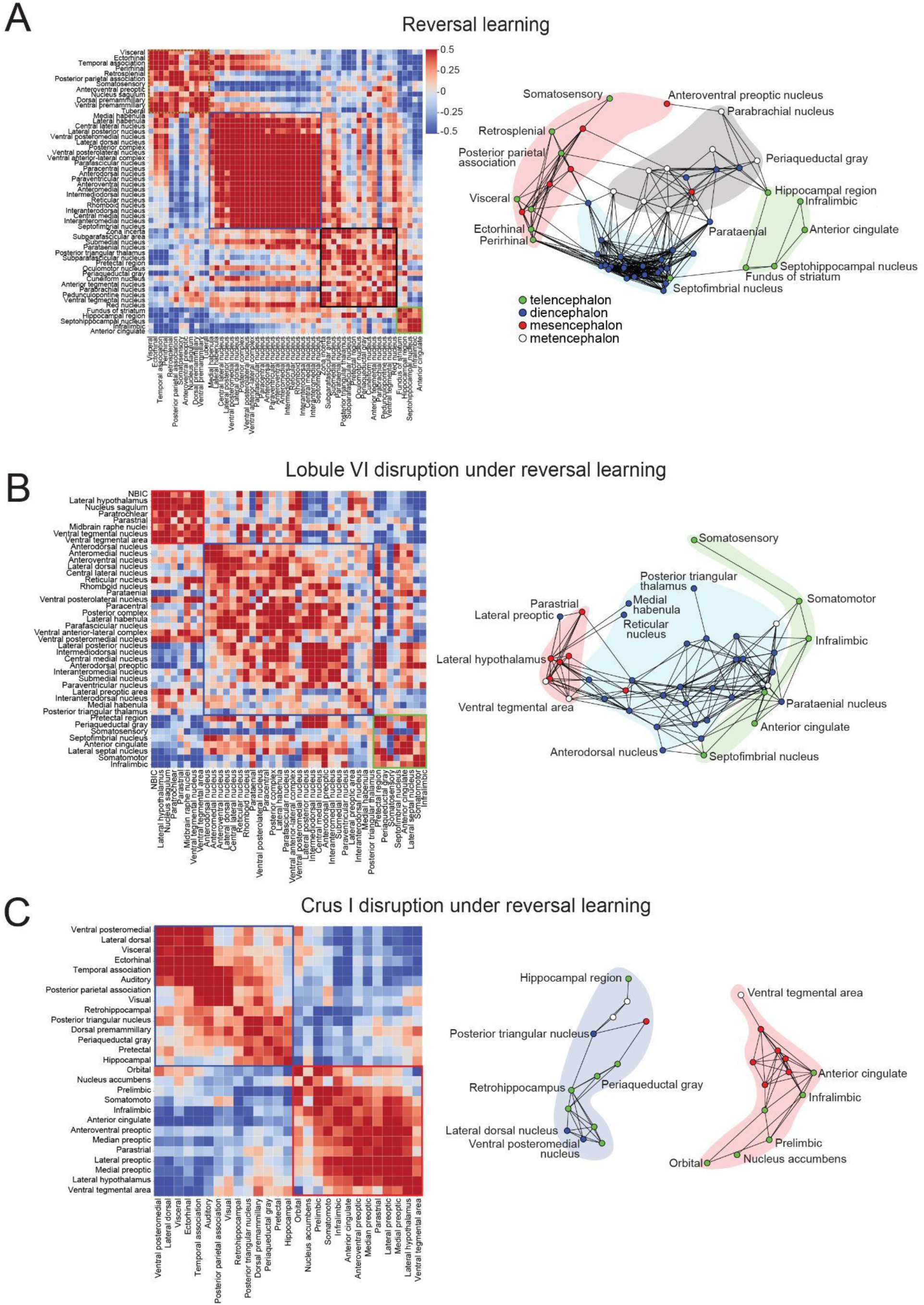
Cerebellar influence on Y-maze reversal networks. Correlation matrices of significant brain regions (p < 0.05) showing inter-region connections for c-Fos expression in (A: left) CNO only reversal, (B: left) lobule VI disruption, and (C: left) crus I disruption (all crus groups combined). Strength of correlation reflected in scale bar (Spearman’s р). Network graphs were generated based on correlations (Spearman’s |r|>0.4) in (A: right) CNO only reversal, (B: right) lobule VI disruption, and (C: right) crus I disruption to visualize relationships between major brain structures (mesencephalon: red, telencephalon: green, metencephalon: white, diencephalon: blue). Lobule VI and crus I inhibition reduces thalamic connectivity required by Y- maze reversal. Abbreviation: Nucleus of the brachium of the inferior colliculus (NBIC).

In reversal learning, we found multiple groups of regions within which correlations were strong (ρ = 0.5-1) (Fig 6A). Diencephalic regions, which encompassed thalamic nuclei, lateral habenula, and paraventricular nucleus, showed the strongest internal correlations. This activity was generally anticorrelated with activity in neocortical regions. Neocortical regions were separated into two groups that were not joined to one another by strong pairwise correlations. One group included retrosplenial, ectorhinal, and retrosplenial cortex, and the other group included anterior cingulate and infralimbic cortex. Inhibition of lobule VI reduced the degree of correlation among thalamic nuclei, and showed strong correlations within mesencephalon, and weak correlations within neocortex (Fig 6B). Within the all-crus-I group, which contained largely mesencephalic and telencephalic regions, within-neocortex correlations formed two groups with no mutual correlations: one including frontal regions (prelimbic, orbital, infralimbic and somatomotor areas, as well as nucleus accumbens) and another for spatial orienting and memory (ectorhinal area, temporal association area, and hippocampal regions) (Fig 6C). In summary, we found that regions specifically activated during Y-maze reversal learning formed thalamocortical functional networks that were disrupted in different patterns after perturbation of lobule VI and crus I.

### Lobule VI and crus I modulate multiday behavioral adaptation to an open field

To test whether the consequences of cerebellar perturbation extended to more complex behavior, we measured the capacity of mice to adapt to a novel environment. We characterized spontaneous behavior in an open-field arena [42] recording mouse behavior by imaging from beneath for 20 minutes over two days in order to track location and to allow automated tracking of body parts using the LEAP (<u>L</u>EAP <u>E</u>stimates <u>A</u>nimal <u>P</u>ose) software package [43], a neural network trained to track the positions of 18 joints (Fig 7A-B **and S3 Movie**).

**Fig 7.**
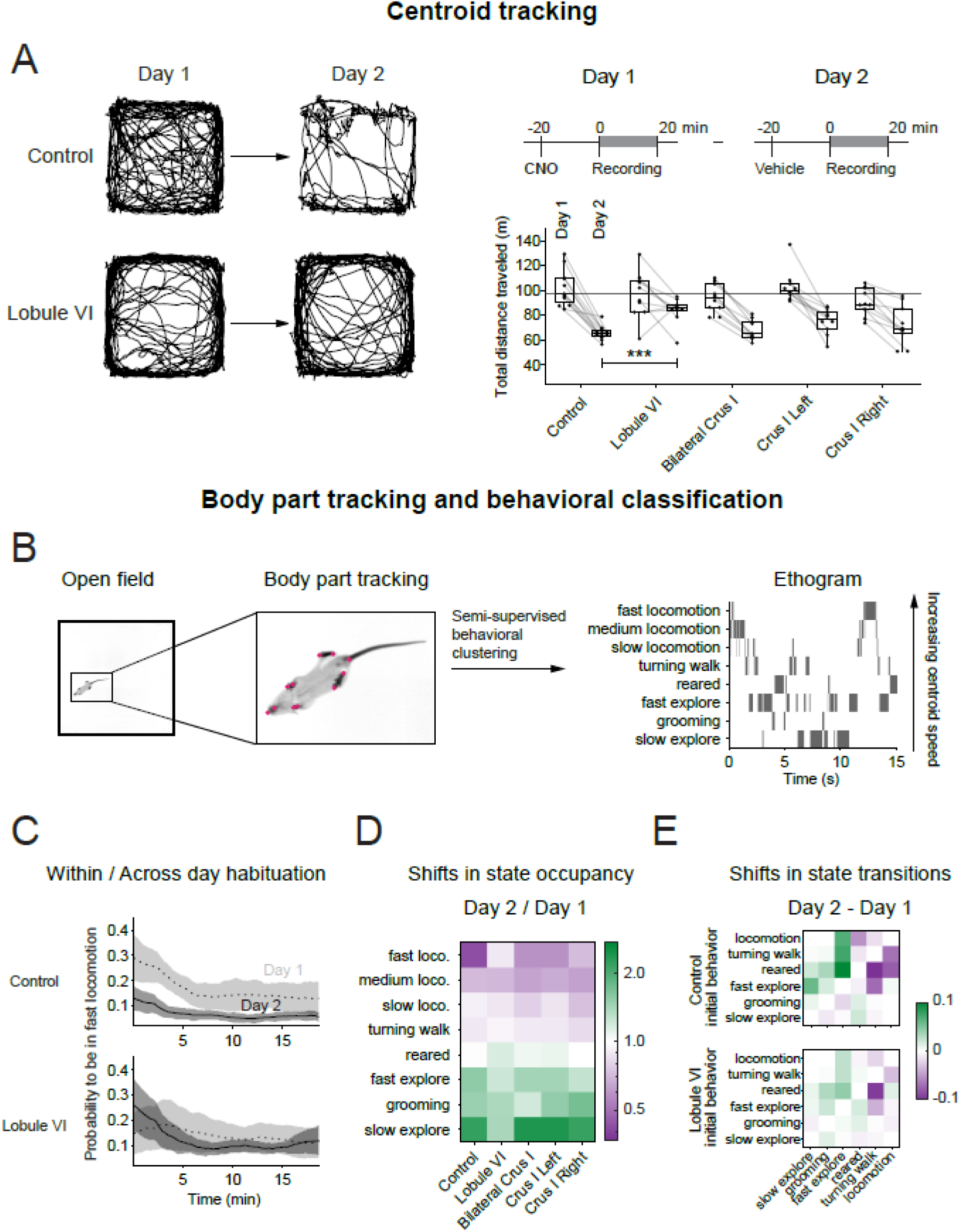
Effects on spontaneous behavior after perturbation of lobule VI and crus I. (A) Lobule VI-perturbed mice show a larger total distance traveled on the second day compared to a day-matched CNO only control group. Left: Example trajectories of the centroid position of a lobule VI-perturbed mouse and a control mouse in the open field for the first 5 minutes. Top right: Timeline of open-field experiments. Comparisons were made using an one-way or two-way mixed ANOVA. (B) Machine learning pipeline to obtain ethograms for the open-field recordings. First, body parts were tracked using LEAP. These postures were processed, as described in Klibaite et al. 2022, to assign one of six behaviors to each time point in the recording. Locomotion was divided into slow, medium and fast based on the centroid velocity of the mouse. (C) The probability to be in the fast locomotion state decreased within and across days for the CNO only control group, but lobule VI-perturbed mice show a higher or similar probability to be in fast locomotion at the beginning of the experiment on the second day compared to the first day. (D) The state occupancies for lobule VI-perturbed mice changed less across days than the other groups, indicating a lack of adaptation. (E) The state transitions for lobule VI-perturbed mice changed less across days than control group, indicating a lack of adaptation. *** p < 0.001

We recorded animals for 20 minutes on each of two days, after a dose of CNO on day 1 and a dose of vehicle on day 2. Over successive days, control mice that did not receive AAV reduced their daily amount of locomotion (two-way mixed ANOVA F(1,47) = 146, p < 0.001). Disruption of lobule VI activity prevented this adaptation. Lobule VI-perturbed mice traveled significantly more on the second day (*d* = 2.3, one-way ANOVA F(4,47) = 5.04, p = 0.002, Dunnett post-hoc test p < 0.001) compared to the control group. The total distance traveled on day 2 was not significantly different from the distance on day 1 in lobule VI perturbed mice, in contrast to a decrease for mice with a unilateral or bilateral disruption of crus I (*d* = 2.6, 2.4, and 1.4 for bilateral crus I, left crus I, and right crus I), or control mice (*d* = 3.4, repeated-measures ANOVA for each group, p < 0.01, Bonferroni correction) (Fig 7A).

Perturbation of lobule VI or crus I did not change the fraction of time mice spent in the inner region of the open field arena compared to control animals on day 1. Right crus I-perturbed animals did spend a significantly larger fraction of time in the inner region of the open field arena on day 1 (r = 0.6, Kruskal-Wallis test chi-squared(4) = 9.52, p = 0.049, pairwise comparisons using Wilcoxon rank sum exact test p = 0.02, Benjamini-Hochberg correction) (**S13A Fig**). None of the manipulations affected locomotory gait (**S13B-C Fig**) or spatial preferences in the arena such as locomotion in the periphery and grooming in corners (**S13D Fig**).

To explore the structure of this altered behavior, we used automated pose analysis. We performed semi-supervised behavioral clustering on LEAP-tracked body-part locations to identify six clusters of body dynamics: slow exploration, grooming, fast exploration, rearing, turning walk, and locomotion. We took the clusters as behavioral states to generate an ethogram, arranged in order of increasing centroid speed (Fig 7B). Since mice spent the most time in locomotion, we further subdivided locomotion into three groups, slow, medium, and fast, based on centroid velocity (**S13C Fig**).

The fractions of time spent in each of the eight behaviors was significantly different for lobule VI-perturbed mice compared to the control animals. More specifically, compared with control animals, lobule VI disruption led to decreases in time spent in the rearing state on day 1 and increases in fast locomotion on day 2 (Fig 7D**, S14A-B,D Fig**). The ratio of fast locomotion in lobule VI-perturbed mice compared to the control group on day 2 was 1.7, sufficient to account for the increased total distance traveled on day 2 compared to the other groups. Within each day’s 20 minutes of observation, the probability of being in the fast locomotion state decayed over time. However, lobule VI-targeted, bilateral crus I-targeted, and right crus I-targeted mice were more likely to perform fast locomotion just after the experiment started on day 2 compared to day 1, in contrast to our observations for the control group and left crus I perturbation (Fig 7C **and S14B Fig**).

We calculated complex adaptation as the ratio of time spent in each state for day 2 compared to day 1. There were few differences among the various controls, except that compared with vehicle-only mice, CNO-exposed animals had a slight reduction in locomotion and more adaptation (ratio further from 1) for fast locomotion and turning walk (**S14C-D Fig**). Overall, the adaptation ratio was closer to 1 for lobule VI-targeted mice for most behavioral states, especially fast locomotion and slow exploration (Fig 7D **and S14C Fig**). We next examined transition probabilities between behavioral states. In control animals, day 2 probabilities compared to day 1 showed higher transition frequencies in the direction of less-active states (*i.e.,* above the diagonal of the matrix in Fig 7E). This tendency was markedly reduced in lobule VI-targeted animals. In crus-I targeted mice, the same semi-supervised behavioral clustering analysis found subtle differences, in particular a right crus I-induced shift from fast-locomotion state to the slow-locomotion state starting on day 1 (Fig 7D **and S14C Fig**). In summary, lobule VI-perturbed animals maintained similar within-day response patterns to the same environment despite impaired adaptation over two days of exposure.

## Discussion

We found that intact cerebellar function in lobule VI and crus I was necessary for two acquired flexible behaviors, choice reversal in a swimming Y-maze and adaptation to an open field. In both behavioral paradigms, silencing of Purkinje cells led to deficits that became apparent over a period of several days, and in the case of Y-maze was marked by incorrect repetitive choices even in a forced session. Y-maze choice reversal recruited activity in a diencephalon-centered group of regions, most of which were reversed by inhibition of Purkinje cells in lobule VI; and in prefrontal and hypothalamic regions, which were reversed by inhibition of either lobule VI or crus

1. Taken together, these studies comprise a demonstration of cerebellar perturbation leading to specific alterations in whole-brain activity and nonmotor function.

Identification of effects on flexible behavior required us to distinguish them from changes in the coordination of movement. In the case of Y-maze reversal learning, we did this using distance swum, a traditional measure of movement. In the arena, pose analysis [42] allowed additional detailed analysis in which we could simultaneously analyze the detailed kinematics of limb movement and longer time-scale features of behavior. Such an analysis required a method that could track individual body parts, such as LEAP [43,4] or other approaches [45,6]; [47,8]. We did not find differences in gait or in spatial occupancy of the arena, suggesting that the chosen cerebellar perturbations affected the evolution of motor behaviors over several days, but not the capacity to interact with the physical environment or generate locomotor behavior. These results are consistent with past work in which rodent gait was not altered by lobule-specific perturbation of posterior cerebellum [16,8], but was changed by cerebellum-wide disruption [42,49,50].

Flexible cognition, as examined by Y-maze reversal learning, was found to be strongly modulated by lobule VI and crus I. Mice demonstrated perseveration in this task by swimming repeatedly toward the previously learned arm before finding the platform in the third arm of the Y-maze, even when the incorrect arm was blocked. Perseverative behavior is a principal criterion for autism spectrum disorder [51–3]. Vermal lobules VI-VII are altered in their volume developmental trajectory in ASD children compared with the rest of cerebellum [14,5], suggesting a specific role for lobule VI in driving ASD-like outcomes. In the open field, lobule VI- perturbed mice demonstrated reduced adaptation by staying in a fast locomotive state on both days. Given the widespread disruption of thalamic activity caused by inhibiting lobule VI output in mice, these behavioral deficits may arise from the disruption of sensory or polymodal processing, reducing the capacity of the forebrain to detect novelty or process its consequences.

We found that disruption of crus I altered both multiday adaptation to the open field and the ability to reverse learning in a swimming Y-maze. Crus I has been implicated in social processing both in mice and in humans with ASD [18,4] and human crus I is engaged in the processing of sensory novelty [17,55,56]. Crus I in rats and mice is putatively homologous to human crus I/II [17] which indicates that our perturbations span the homologue of human ansiform area. Taken together, our findings are in accordance with lobule-specific substrates for a number of deficits that arise in cerebellar cognitive-affective disorder [6,7,57].

The cerebellum influences the rest of the brain through polysynaptic paths through the deep and vestibular nuclei. By analyzing the immediate-early gene c-Fos, we found that reversal learning engaged midbrain, diencephalic, and neocortical regions associated with flexible behavior, and that this engagement was altered by perturbation of lobule VI or crus I. Many of these brain regions receive disynaptic paths from lobule VI and crus I, as demonstrated by transsynaptic tracing [1–5]. In the mesencephalon, the habenula, periaqueductal gray, and parabrachial nucleus are engaged during defensive, negative-reward, and decision-making behavior [58–1]. Finally, the infralimbic and anterior cingulate cortex are activated in effective decision making and reward-seeking [62–4]. Our observation that lobule VI and crus I inhibition affects activity in these regions during reversal learning suggests that cerebellar activity is necessary for the normal expression of a wide range of brain activity in the face of changing environmental valence.

Recent studies in mice show that vermal and hemispheric regions project via the deep cerebellar nuclei to distinctive patterns of forebrain structures [1–4,65,66], influencing thalamocortical nonmotor processing. Chemogenetic inhibition of lobule VI Purkinje cells led to broad decreases in thalamic activity as measured using c-Fos. Inhibition of lobule VI also led to a strong loss of functional network correlation among thalamic structures, as well as among neocortical regions. A major target of cerebellar output, especially from lobule VI/VII, is the thalamic reticular nucleus [1–5], which is inhibitory and sends its outputs throughout the rest of the thalamus. Reticular nucleus paths have been suggested to have a gating effect on thalamocortical function [67] and are important for flexible behavior [68–1]. In addition, removal of Purkinje-cell inhibition in lobule VI might be expected to increase activity in thalamic polymodal nuclei via cerebellothalamic excitation. The fastigial nucleus, which receives strong lobule VI input, has recently been shown to send output to brainstem targets subserving arousal and autonomic functions [66]. The widespread nature of lobule VI’s functional targets supports the idea that the cerebellum acts as a powerful modulator of thalamocortical processing during flexible behavior. Our findings suggest that lobule VI pathways are also necessary for coordination among thalamic regions.

Chemogenetic inhibition of crus I Purkinje cells also led to changes in neocortical activity as measured using c-Fos. Human and rodent crus I [1,2,4,5,72] project to thalamus, both sensory/motor such as ventral posteromedial nucleus and polymodal such as lateral dorsal nucleus, and to hypothalamus, which like the thalamus is a diencephalic structure that projects monosynaptically to neocortex [73]. Transsynaptic paths from rodent crus I project particularly densely to infralimbic, prelimbic, and orbital cortex, providing a substrate for our observed alterations in c-Fos expression [5]. We found that under crus I inhibition, activity in neocortical regions differed strongly from unperturbed reversal learning, and functional network analysis revealed separation of activity into highly distinct sensory and associative neocortical networks, suggesting that during flexible Y-maze learning, crus I plays a role in coordinating the action of these two networks.

Our c-Fos results could have potentially arisen from either direct consequences of cerebellar perturbations, or indirect long-term consequences arising after failure to complete the task. Two lines of evidence suggest that at least some effects were direct consequences. First, lobule VI perturbation changed thalamic c-Fos activation more than it changed neocortex, and crus I perturbation changed neocortex but not thalamus. This difference is consistent with the observation that neocortex receives inputs from many nonthalamic sources, and indeed disjoint activation of thalamus and neocortex has been long known [74,5]. Second, differences in total distance swum were considerably smaller than the observed changes in c-Fos expression. It is nonetheless true that c-Fos expression depends in a complex and cell-type-dependent manner on activity [76] and persists for many minutes, far longer than the timescale of synaptic chains of activation (<1 s). Our work identifies candidate regions for future electrophysiological recording, which can unambiguously identify the direct effects of cerebellar perturbation.

Lateralization of cerebellar function has previously been reported in mice for social interaction and grooming [18,4]. We observed that right crus I-targeted perturbation affected open-field adaptation. Previous work found that injections targeted to either left or right crus I alone were sufficient to impair reversal learning [18], but we found that perturbation of both sides was needed to cause impairment. These differences could be explained if our injections had a different amount of spillover outside targeted regions than previous work. Our observation that targeted injection can cause spatially distributed expression highlights the need to quantify the spatial range of DREADD-based perturbation.

In summary, we have used chemogenetic inhibition of Purkinje cells to identify two cerebellar regions that influence multiday flexible behavior, lobule VI and crus I. Detailed characterization of c-Fos activation in Y-maze, including different stages of learning (habituation, acquisition/“nonreversal”, and reversal), revealed brainwide consequences of the task and focal perturbations. First, more challenging task conditions led to progressively more widespread regional activation. Second, cerebellar perturbation affected activity in regions that were activated under reversal learning, including thalamus by lobule VI and neocortex by crus I. Brain-wide c-Fos mapping serves as a screen analogous to the introduction of the Genome-Wide Association Study in 2002 [77]. Understanding the task-specific role for cerebellum in driving forebrain neural processing will require direct recording or high-time-resolution perturbation of the candidate regions we have identified.

## Methods

### Experimental Design

We targeted neural activity of Purkinje cells of mice using inhibitory DREADDs (Fig 1A**, S1A-C Fig)**. Mice used in this study were male C57BL/6J (The Jackson Laboratory, Bar Harbor, ME) and acclimated for at least 48 hours at the Princeton Neuroscience Institute vivarium prior to procedures. After a three-week recovery period, mice (PND 70) underwent behavioral testing starting with open field and thenwater Y-maze reversal for whole-brain c-Fos analysis (Fig 1I; Table 1).

All mice were housed in Optimice cages (Animal Care Systems, Centennial, CO) and received environmental enrichment, including paper nesting strips and one heat-dried virgin pulp cardboard hut (Shepherd Speciality Papers, Milford, NJ). Mice were fed PicoLab Rodent Diet food pellets (LabDiet, St. Louis, MO) and water was provided ad libitum. Animals were housed in groups of 4-5 mice in reverse light cycle rooms to maximize normal nocturnal activity, as behavior testing occurred during the day. All experimental procedures were approved by the Princeton University Institutional Animal Care and Use Committee and in accordance with animal welfare guidelines of the National Institutes of Health.

### Animal preparation

We delivered adeno-associated virus (AAV) with the sequence for the inhibitory DREADD hM4Di, which was fused to mCherry protein under a Ef1α promoter. This virus included a DIO component, which when combined with the L7-cre virus, expressed in the Purkinje cell layer exclusively (Fig 1A**; and S1A-C Fig**). Recently clozapine-N-oxide (CNO) has been found to convert back to the parent compound, Clozapine, in mice prior to crossing the blood-brain barrier. To reduce confounds in our experimental design, all mice received CNO during all behavioral tasks, as Clozapine may alter signaling of neuromodulators, notably dopamine and serotonin[78–80].

For DREADD electrophysiology and behavioral experiments, lobules targeted were lobule VI (n = 36 behavior, n = 10 for recordings), bilateral crus I (n = 17), crus I right (n = 25), crus I left (n = 26), but as AAV can spread into neighboring lobules, mCherry fluorescence was recovered and quantified for the entire cerebellum. Controls included animals injected with AAV without DREADDs (CNO and mCherry, n = 18), CNO only (n = 30), Vehicle (DMSO and saline, n = 9), and untreated (n = 123). To understand if CNO or a lobule-specific perturbation altered Y-maze performance without reversal a subset of CNO only (n = 7) and CNO and Lobule VI mice (n = 10) underwent 25 trials of a third day of acquisition. To understand learning in the Y-maze, animals were sacrificed after habituation (n = 16), acquisition day 1 (n = 16), and acquisition day 2 (n = 10) for a total of 42 mice. When conditions allowed, the same animals went through both water Y-maze and open field testing, see Table 1.

Briefly, animals were anesthetized with isofluorane (5% induction, 1-2% oxygen; 1 L/min) and mounted in a stereotaxic device (David Kopf Instrument, Tujunga, CA) for all surgeries. Temperature was monitored and automatically adjusted using PhysioSuite (Kent Scientific Corporation, Torrington, CT). Animals were prepared for surgery with an application or Puralube vet ointment (Pharmaderm Florham Park, NJ) to prevent corneal drying, the scalp was shaved and cleaned, and animals received osmotic diuretic drug 15% D-mannitol in DPBS (0.02ml/g; intraperitoneal, i.p.) and an anti-inflammatory drug, Rimadyl (5mg/kg Carprofen 50 mg/ml, Pfizer, Eurovet, in NaCL; every 24 hours for 2 days; subcutaneous, s.c.). A lateral skin incision was made over the lambdoid suture. Muscle was cut over the occipital bones first vertically than horizontally and as close to the bone as possible to allow for regrowth post-surgery and enough to expose lobule VI or crus I. A small craniotomy was made over each lobule of interest for injection of inhibitory DREADD AAV1-Eflɑ-DIO-hM4D(Gi)-mCherry-WPRE-hGHpA (8.5 x 10^13; PNI Vector Core, AAV-VC68) or control AAV8-Eflɑ-DIO-mCherry-WPRE-hGHpA (1 x 10^15; PNI Vector Core, AAV-VC139). To target Purkinje cells, both DREADD and control AAVs were mixed in a 1:1 ratio with AAV1-sL7-Cre-HA-WPRE-hGH-pA (2 x 10^14; PNI Vector Core, AAV-VC141). Virus was injected using borosilicate glass capillaries (World Precision Instrument, Sarasota FL) made using the Sutter Micropipette Puller (Model P-2000, Sutter Instrument Company) and bevelled at a 45 degree angle. To ensure viral spread ∼600nl total of DREADD or control was injected per mouse, distributed at 3 separate depths (150, 250, 450 µm below the dura) and two locations per lobule. Craniotomy was sealed with a silicone elastomer adhesive (Kwik-Sil, World Precision Instrument, Sarasota, Fl) and skin was sutured.

### Acute brain slice experiments

Mice (C57BL/6J) were prepared as described previously in “Animal preparation” for inhibitory DREADD induction of Purkinje cells at 3-weeks of age (n = 4). Two weeks later, mice were deeply anesthetized with Euthasol (0.06 ml/30g), decapitated, and the brain removed. The isolated whole brains were immersed in ice-cold carbogenated NMDG ACSF solution (92 mM N-methyl D-glucamine, 2.5 mM KCl, 1.25 mM NaH_2_PO_4_, 30 mM NaHCO_3_, 20 mM HEPES, 25 mM glucose, 2 mM thiourea, 5 mM Na-ascorbate, 3 mM Na-pyruvate, 0.5 mM CaCl_2_, 10 mM MgSO_4_, and 12 mM N-acetyl-L-cysteine, pH adjusted to 7.3–7.4). Parasagittal cerebellar brain slices (270 μm) were cut using a vibratome (VT1200s, Leica Microsystems, Wetzlar, Germany), incubated in NMDG ACSF at 34°C for 15 minutes, and transferred into a holding solution of HEPES ACSF (92 mM NaCl, 2.5 mM KCl, 1.25 mM NaH_2_PO_4_, 30 mM NaHCO_3_, 20 mM HEPES, 25 mM glucose, 2 mM thiourea, 5 mM Na-ascorbate, 3 mM Na-pyruvate, 2 mM CaCl_2_, 2 mM MgSO_4_ and 12 mM N-acetyl-L-cysteine, bubbled at RT with 95% O_2_ and 5% CO_2_). During recordings, slices were perfused at a flow rate of 4–5 ml/min with a recording ACSF solution (120 mM NaCl, 3.5 mM KCl, 1.25 mM NaH_2_PO_4_, 26 mM NaHCO_3_, 1.3 mM MgCl_2_, 2 mM CaCl_2_ and 11 mM D-glucose) and continuously bubbled with 95% O_2_ and 5% CO_2_.

Whole-cell recordings were performed using a Multiclamp 700B (Molecular Devices, Sunnyvale, CA) using pipettes with a resistance of 3–5 MΩ filled with a potassium-based internal solution (120 mM potassium gluconate, 0.2 mM EGTA, 10 mM HEPES, 5 mM NaCl, 1 mM MgCl_2_, 2 mM Mg-ATP and 0.3 mM Na-GTP, pH adjusted to 7.2 with KOH). Purkinje neurons expressing mCherry were selected for recordings (Fig 1C-D).

### In vivo electrophysiological recordings

Two days after performing craniotomy and headplate implantation, animals (n = 6) were habituated to head fixation on the treadmill. The day of the recording, cover was removed from the cranial window and then, a Neuropixels probe 1.0 (Imec, Belgium)[81] coated with a fluorescent dye (CM-DiI, Thermofisher) was slowly inserted (0.1 mm per 1 minute) using motorized micromanipulator (MP-225; Sutter Instrument Co.) into the cerebellum with the tip reaching depth of 3-5.5 mm below the brain surface. Once the right depth was reached, the probe was left to rest for 15-30 min, before starting the recording. Sterile saline was used to cover the exposed cerebellum. Signals from 384 electrodes were recorded simultaneously at 30 KHz using the Neuropixels headstage 1.0 and Neuropixels 1.0 PXIe acquisition system (Imec). High-frequencies (>300 Hz) and low-frequencies (<300 Hz) were acquired separately. SpikeGLX software (http://billkarsh.github.io/SpikeGLX/) was used to select the recording electrodes, adjust gain corrections and save data. Tactile sensory stimulation was performed in awake mice using the air puffs (40 ms, 20 psi, randomized inter-trial interval, 100 trials) delivered ipsilateral to the recording site via a small tube (2 mm diameter), approximately placed parallel to the anterior-posterior axis, 10 mm mediolateral and 1 mm anterior to the nose of the mouse, and connected to solenoid valve (The Lee Co.) controlled by paired microcontrollers (Arduino Due) and a single board computer (Raspberry Pi). Timings of air puff stimulation were digitized at 10 kHz with multifunction DAQ module (PXIe-6341 unit with BNC-2110 breakout box, National Instruments) and synchronized with using TTL pulses from PXIe acquisition module. Spikes were sorted offline using Kilosort2[82], using default parameters. Manual curation of clusters were performed using Phy (https://github.com/cortex-lab/phy). After extracting timestamps of each putative single unit activity, peristimulus time histograms and firing rate changes were analyzed and plotted using a custom MATLAB script. All recording sites were confirmed by post-hoc histology, in 100 µm coronal cerebellar sections recording tracks were identified with CM-DiI marks (Fig 1E).

### Tissue processing and histology

To examine mCherry fluorescence, the presence of DREADD virus, two mice were anesthetized with Euthasol (0.06 ml/30g, i.p.) and perfused with 4% paraformaldehyde (PFA). Brains were stored overnight at 4% PFA then placed in 20% sucrose in PBS overnight until sectioning at 50 µm. Sections were washed with PBS and incubated for 1 hr at room temperature (RT) in blocking buffer (10% normal donkey serum, 0.5% Triton in PBS) prior to an overnight incubation at 4℃ in PBS buffer with 2% normal donkey serum, 0.4% Triton and the rabbit anti-RFP (600-401-379, Rockland Immunochemicals, Inc., Limerick, PA; 1:1000) primary antibody. The next day, sections were washed in PBS and incubated for 2 hr at RT in PBS buffer with 2% normal donkey serum, 0.4% Triton, and the donkey anti-Rabbit Alexa Fluor 647-conjugated secondary antibody (A-21449; Thermo Fisher Scientific, MA, USA, Invitrogen; 1:400).Tissue was mounted on glass slides with Prolong Diamond (ThermoFisher Scientific, MA, USA). Tissue was imaged with a Leica SP8 confocal laser-scanning microscope (Leica Microsystems, Germany) using 40x or 63x objectives (**S1B-C Fig**).

To confirm tissue clearing and lightsheet microscopy, brains were sectioned and stained using conventional immunohistochemistry. Free-floating tissue sections were cut using a vibratome (VT-1000S, Leica Microsystems, Germany). After washing in phosphate buffered saline (PBS) ph 7.6 at RT for four 10 min sessions, sections were blocked with 10% donkey serum and 0.5% Triton X-100 in PBS for 1 hr. Post-incubation, sections were placed in the primary solution of rabbit anti-Fos (226 003, Synaptic Systems,Goettingen, Germany, 1:1000) with 2% donkey serum and 0.5% Triton X-100 in PBC for 48 hours at 4°C. Sections were washed for three 10 min sessions in PBS then placed in secondary donkey anti-Rabbit Alexa Fluor 647-conjugated secondary antibody (A-21449; Thermo Fisher Scientific, MA, USA, Invitrogen; 1:200) in 2% donkey serum and 0.4% Triton X-100 in PBS for 2 hr at RT. Sections were washed for three 10 min sessions in PBS. Tissue was imaged at 20x using an epifluorescence microscope (NanoZoomer S60, Hamamatsu Photonics) and c-Fos positive cells were counted in ImageJ (**S7A and D Fig**).

After animals completed the last Y-maze reversal session, animals were placed back in their home cage for 90 minutes. Then mice were anesthetized with Euthasol (0.06 ml/30g, i.p.) and perfused with 4% PFA for analysis of c-Fos and mCherry expression for DREADD recovery using iDISCO+ clearing methods[5,35,36]. Briefly, after an overnight fix in 4% PFA, brains were rinsed in PBS at RT for four 30 minute sessions. Immediately brains were dehydrated 1 hr at each ascending concentration of methanol (20, 40, 60, 80, 100, 100%) and placed overnight in 5% hydrogen peroxide and methanol at RT. The next day, brains were rehydrated for 1 hr at each descending concentration of methanol (100, 80, 60, 40, 20%) and lastly PBS. Samples were placed in detergent (0.2% Triton X-100 in PBS) for two 1 hr sessions then placed for two days in 20% Dimethyl sulfoxide (DMSO), 0.3 M glycine, 0.2% Triton X-100 in PBS at 37℃.

Brains were blocked in 10% DMSO, 6% donkey serum, 0.2% Triton X-100 in PBS at 37°C for 3 days. Once at RT, samples were washed in PTwh (0.2% Tween-20, 10µg/ml heparin in PBS) and placed in primary solution of rabbit anti-Fos (226 003, Synaptic Systems,Goettingen, Germany, 1:1000) and/or rabbit anti-RFP (600-401-379, Rockland Immunochemicals, Inc., Limerick, PA; 1:450) for one week at 37℃. Brains were washed in PTwH five times in increasing amounts of time (10, 15, 30, 60, 120 min) and then placed in secondary donkey anti-Rabbit Alexa Fluor 647-conjugated secondary antibody (A-21449; Thermo Fisher Scientific, MA, USA, Invitrogen; 1:200) for one week at 37℃. Brains were washed in PTwH five times in increasing amounts of time (10, 15, 30, 60, 120 min) then dehydrated 1 hr at each ascending concentration of methanol (20, 40, 60, 80, 100, 100%) until being placed in 66% DCM/33% methanol for 3 hrs at RT. Brains were cleared with 100% dichloromethane (DCM) for two 15 min steps then placed in 100% benzyl ether (DBE). Brains were kept in fresh DBE prior to imaging on a lightsheet microscope and after for long-term storage.

### Water Y-maze

The water Y-maze assay and analysis was similar to described [16]. Briefly, the Y-shaped transparent polycarbonate apparatus had symmetrical arms, each measuring 33 cm x 7.5 cm x 20.3 cm (length x width x height) from the center. Notches in all three arms (9.5 cm from the center) allowed for a removable gate. A Pyrex glass container was used as a platform for the mice to climb on. To prevent the animals from seeing the platform, opaque water was used by mixing ACMI certified hypoallergenic non-toxic white paint (Art-Minds, Tempera Paint) in warm water. Water levels were kept at a depth to prevent the animals from touching the bottom of the maze. Between each mouse, excrement was removed and water was exchanged to maintain an ideal water temperature between 22-28 degrees Celsius. At the end of the day, water was removed for cleaning by PREempt disinfectant wipes (0.5% Hydrogen Peroxide) and sprayed down with 70% ethanol to be left overnight to dry. To prevent distraction, a three-walled shield was placed around the maze constructed of non-reflective black plastic (34 x 29 x 22 cm, length x width x height). Directly above the arena, a camera (PlayStation Eye) was mounted on a T-slotted aluminum rail (80/20 KNOTTS Company, Berkeley Heights, NJ) and used to record the entire field of view at 50 frames/s using a custom Python 2.7.6 script (Anaconda 1.8.0) and a CLEye Driver (https://codelaboratories.com/products/eye/driver/).

Over four consecutive days animals were habituated to the arena (Day 1; three 60 s trials), learned to find a platform through trial and error (Acquisition Day 2 and Day 3; four sessions with five 40 s trials), then exposed to reversal whereby the platform was moved to the opposite arm the animals learned (Day 4; five sessions with five 40 s trials) (Fig 2A). Mice were required to have 80% success rate for acquisition day 3 in order to progress to the reversal day. During the reversal day, animals were exposed to four sessions of five consecutive trials followed by a fifth forced session whereby a door was placed in the initial learned arm of the maze which no longer has a platform. All mice were kept in a clean cage on a heating pad to dry before returning to their homecage. As the temperature difference between the warming cage and the water can be drastic, it is critical to allow for the animal to cool down prior to starting a new session. Performance on first-choice turn direction in the Y-maze assay was calculated automatically. Neither surgery by itself, nor the effects of administering vehicle-only, CNO-without-DREADD, or CNO-with-mCherry, affected distance swum, initial learning, or reversal-learning compared with untreated mice (**S3A-C Fig**). Distance traveled during habituation was calculated to assess possible muscle damage during surgery (**S1A Fig**). Subsequent tries were recorded to calculate the fraction of choices required to reach the platform (Figure 2D **and S3F Fig**).

### Lightsheet microscopy

Briefly, cleared brains were glued (Loctite, 234796) ventral-side-down to a 3D printed holder and imaged in DBE. Brains were registered using the autofluorescence channel (488 nm laser diode excitation and 525 emission) to the Princeton Mouse Atlas. Cellular imaging of c-Fos and mCherry expression was acquired using 640 nm excitation and 680 nm emission (1x magnification, 1.3x objective, 0.1 numerical aperture, 9.0 mm working distance, 12.0 x 12.0 mm field of view, LVMI-Fluor 1.3x, LaVision Biotech) with a 10 μm step-size using a 0.016 excitation NA. Analysis of whole-brain c-Fos was completed using ClearMap[37] on high performance computing clusters. To confirm ClearMap cell counts, two human annotators analyzed 14 brain volumes and found 96.72% correspondence between cells counted by the human annotators and ClearMap counted cells. Structures less than 80 microns and structures in the medulla were removed from analysis. The medulla was not analyzed as it can be damaged during brain extraction. Ventricles, brain edges, and zones within 60 microns of region boundaries were removed. The cerebellum was not analyzed for c-Fos activity. Visualization of brain volumes and cell detections from ClearMap was performed using Neuroglancer. Tissue image processing and registration was performed using custom Python code (https://github.com/PrincetonUniversity/BrainPipe).

We assessed the impact of varying experimental conditions in the following contrasts:

1. Acquisition day 1 vs Habituation
2. CNO control reversal vs CNO control no reversal
3. CNO control reversal vs Vehicle control reversal
4. Vector control reversal vs Vehicle control reversal
5. Lobule VI reversal vs CNO control reversal
6. Crus I left vs CNO control reversal
7. Crus I right vs CNO control reversal
8. Bilateral Crus I vs CNO control reversal
9. Lobule VI reversal vs Lobule VI no reversal

In contrasts 1-5, both the control and the treatment groups of mice were processed in the same batch so as to minimize batch effects. In contrasts 6-7, both control and treatment groups were present in multiple batches, so batch (encoded as a categorical indicator variable) was included as a covariate to adjust for the batch effect. Finally, in contrasts 8-9 control and treatment groups did not overlap within any single batch, impeding the direct adjustment for confounding due to batch effects. However, we were able to adjust for confounding indirectly by using a “bridge variable” strategy inspired by the chain-type experimental design described by Song et al. (2020) [83]. We define a bridge variable as an experimental group found in both the batch containing treatment animals as well as the batch containing control animals. The bridge variables were Lobule VI reversal for contrast 8 and CNO control reversal for contrast 9. Inclusion of the bridge variable along with the batch indicator variables in a regression model makes the treatment versus control contrast statistically identifiable separate from the batch effect, as previously shown[83].

Total counts of active neurons were highly variable between animals even in the same experimental condition. We therefore sought to explain differences in proportion of total counts for each brain region. Given the overdispersed, discrete nature of the data, we used a negative binomial likelihood and performed separate regressions for each brain region and contrast, using the natural log of total counts as an offset. Specifically, for a given contrast, let X_i_=1 if animal i is in the treatment group and X_i_=0 otherwise. Let Z_i_=1 if the animal is in the first batch and Z_i_=0 otherwise. Let A_i_=1 if the animal has the bridge variable condition and A_i_=0 otherwise.Let T_i_ be the total c-Fos counts across all regions in brain i. Let Y_ij_ be the c-Fos count in region j of brain i. The statistical model is then given by

Y_ij_ ∼ negative binomial (μ_ij_,φ_j_)

log(μ_ij_) = β_0j_+X_i_*β_1j_ + ln(T_i_) + Z_i_*β_2j_ + A_i_*β_3j_

where μ_ij_ is the expected value of Y_ij_, φ_j_ is the nuisance shape parameter of the negative binomial distribution, and β_0j_ is an intercept term which captures the fact that some brain regions have a consistently higher or lower activity level across all animals without regard to their experimental condition (for example this could be because some regions have a larger volume than others).

The parameter β_1j_ is of greatest biological interest. It is interpreted as the average change in proportion of c-Fos activity for region j in the treatment group relative to the control group on the logarithmic scale. For example, if β_1j_=1.5, the average c-Fos activity for brain region j is estimated to be exp(1.5)=4.48 times higher in the treatment condition compared to the control condition. If β_1j_= -0.7 (a negative coefficient) average c-Fos activity for brain region j is estimated to be exp(-0.7)=0.5 times lower in the treatment condition compared to the control (ie, the treatment mean is half that of the control mean).

We fit each negative binomial regression using the R package MASS [84]. In addition to maximum likelihood estimates of the regression coefficients such as β_1j_, this package also computes a standard error for each coefficient. From these we obtained effect sizes in the form of Wald z-test statistics (estimated coefficient divided by standard error). Under the null hypothesis that there is no change in the average c-Fos expression between the treatment and control groups (i.e. β_1j_=0), the effect sizes would be expected to follow an asymptotically standard normal distribution. By comparing the computed effect sizes against this null distribution, we obtained p-values. If the p-value was small for a given brain region, it suggested that there was a large difference between the treatment and control groups and the null hypothesis should be rejected for that region. Since a separate statistical test was performed for each brain region, we adjusted raw p-values in each analysis to control the multiple testing false discovery rate (FDR) using the method of Benjamini and Hochberg [85].

In rare cases, numerical errors occurred in the MASS package fitting procedure. This is because estimation of the negative binomial parameters (β_0j_, β_1j_, β_2j_, β_3j_, and φ_j_) requires an iterative optimization that can fail to converge. We determined that in most of these cases, the brain region was small and/or irregularly shaped. We therefore dropped such regions from subsequent analysis.

In contrast 8 only, some observations came from batch 201810_adultacutePC_ymaze_cfos, in which brains had cerebellum excised. As a quality control step, we excluded regions with extremely low counts and low variability across all brains in this batch since this would destabilize the regression fitting procedure. Specifically, for each region we counted the number of brains within the batch having a nonzero count value. If the number of nonzero counts was less than two, we excluded the region.

In addition to analyzing individual brain regions, we also fit negative binomial regressions to examine the effects of experimental perturbations on three composite regions consisting of multiple subregions from our original 122 regions: thalamus, sensory/motor, and polymodal association. For each composite region and experimental contrast, we summed the raw c-Fos cell counts of all constituent subregions within each animal. We then fit negative binomial regressions as previously described, with the natural log of the total counts in the entire brain again used as an offset for each animal. This maintained consistency in the interpretation of the regression coefficients and effect sizes by keeping them on the same (proportional) scale as the original analysis. To confirm our c-Fos statistical results, we ran two sample t-tests for each region per condition comparison, and then checked the validity of the resulting p-values using permutation testing. None of the highly significant regions from the negative binomial regression were missing.

### Open Field

Wild-type (n = 60) [42], mice with a cerebellar perturbation (n = 10 per group) and controls, including CNO only (n = 12), CNO and mCherry (n = 10), and vehicle only (n = 10). Previous data collected [42] using Purkinje-cell specific tuberous sclerosis 1 gene mutation (L7Cre; Tsc1^flox/flox^) (n = 9) was used as a comparison for gait analysis (**S13B Fig).** Animals were placed in an open field arena measuring 45.72 x 45.72 cm (length x width) and 30.48 cm in height with a transparent polycarbonate floor, as previously described [42]. A Point Grey grayscale camera (12-bit grayscale, 1280 x 1024 pixel resolution at 80 Hz) was used to image from below. The soundproof box with ventilation was illuminated with far-red LEDs. To prevent noise disturbance, doors were kept closed during acquisition. Mice received CNO on day 1, if applicable, and vehicle on day 2 to understand how perturbation may alter open field habituation. Each mouse was recorded for 20 minutes before returning to group housing over two days (first 18 min 46 sec are included in the analysis). Raw images were processed to segment the mouse from the background (median filter) to a final video size of 400 x 400 pixels.

### Machine learning

Each frame was aligned for the mouse body axis and body parts were tracked using LEAP (<u>L</u>EAP <u>E</u>stimates <u>A</u>nimal <u>P</u>oses) as previously described[42,3]. The LEAP network was trained on 1000 frames to find 18 body parts. Automatic classification of animal behavior was performed using custom MATLAB and Python scripts as previously described [42]. Briefly, distances between 11 body parts (nose, chin, 4 x paw tip, 4 x paw base, where paw connects to leg, and tail base) were calculated and the dimensionality was reduced by projecting on the first 10 PCA components. A wavelet analysis was performed in the lower dimensional space, followed by k-means clustering (k = 100) of the frequency data to obtain behavioral clusters. The 100 behavioral clusters were manually grouped into 6 behaviors (slow explore, grooming, fast explore, reared, turning walk, locomotion) (Fig 7B). A majority filter with a sliding window of 11 frames was used on the predicted behaviors for each frame. Centroid metrics were used to calculate distance traveled, and fraction of time in the inner arena. The inner part of the arena was defined as the region of the arena with a distance larger than 150 pixels (approximately 7.6 cm) to all borders of the arena. The borders were obtained using the space explored by the mouse. To calculate the temporal change of the probability to be in fast locomotion within the time of an experiment, a sliding window of size 20001 frames was used. The probabilities at equally distant time points were calculated for each mouse separately, based on which the mean and standard deviation was obtained (Fig 7C).

For the shifts in state occupancy, the fractions of time spent in each behavior during the first five minutes and the remainder of the experiment were calculated for each mouse. Then, the geometric mean was taken for each group and day and the geometric mean values for day 2 were divided by the ones from day 1 for each group (Fig 7D). The shifts in state transitions were obtained as follows: First, the transition matrices were calculated for each mouse and day by considering only changes of states as transitions (i.e. the transition rate to stay in the same state is zero). Then, the transition matrices are averaged for each group and day by weighting the transition probabilities from initial state i by the probability of the mouse to be in state i. Finally, the resulting averaged transition matrix for day 1 is subtracted from the one for day 2 for each group, which allows an investigation of the changes in state transitions across days. To study the possibility of gait changes in cerebellar perturbed mice compared to controls, the phases when the paws enter stance during a single stride of locomotion were calculated for different centroid velocities[49]. The beginning of the stance phase was determined by the peak position of the paw in an animal-centered coordinate system. All analyses of the open field behavioral data were performed using custom Matlab code (https://github.com/PrincetonUniversity/OF-ymaze-cfos-analysis).

### Statistics

All statistics were performed using MATLAB, R (rstatix, compositions, npmv, data.table, plyr, ggplot2, ggpubr, car, DescTools), or Python 2 and Python 3 (statsmodels, scipy, matplotlib, numpy). Data are presented as mean ± SD, unless otherwise stated. Group mean comparisons were performed through repeated measures ANOVA (open field) or through one-way ANOVAs with Tukey HSD or Dunnett multiple comparisons post-hoc tests. For each comparison the effect size (Cohen’s d) or Kolmogorov-Smirnov test was calculated. Normality was tested using the Shapiro-Wilk test, and the homogeneity of variances was tested using Levene’s test to determine parametric or nonparametric analysis. If the Shapiro-Wilk test or the Levene’s test was significant, the nonparametric Kruskal-Wallis test was performed, followed by pairwise comparisons with Wilcoxon rank sum exact tests (with Benjamini-Hochberg multiple comparison correction). In this case, the Wilcoxon effect size r was calculated for each comparison.

Y-maze performance was measured by the number of successes and failures of each mouse in 5 trials for different sessions and days. To account for the fact that the data were nested and can take only values between 0 and 100% (more specifically 0, 20, 40, 60, 80 and 100%), we fit generalized mixed effect models (GLMM) with a binomial distribution and logit link function to the performance data for acquisition day 2, 3 and reversal day (using glmer function in R package lme4 [86]. We included the session and the experimental group as fixed effects predictors. Since the performance scores within mice may be correlated, we also incorporated random intercepts in the model. We tested different models (with and without the interaction term of group and session; considering session as a factor or a quantitative variable; with and without random slopes added to random intercepts) and chose the models based on the Akaike information criterion (AIC) and likelihood ratio tests. The data for acquisition day 2 and day 3 was best described by a model without the interaction term of group and session and random intercepts only. To test for significant differences of the performance of the experimental groups, we performed multiple comparisons of the means (Dunnett contrasts, using the glht function in the R package multcomp) [87]. For the reversal day, we also considered a model that accounts for interactions of group and session and tested for within session differences between the groups (Bonferroni-Holm multiple comparison correction).

The number of structures used for c-Fos analysis were tested for multiple comparisons by calculating the false discovery rate, the coefficient of variation, and by performing permutation tests on each comparison. All c-Fos data is presented as p<0.01 unless otherwise stated. To analyze correlations between whole-brain c-Fos/DREADD and behavior, Spearman’s rank correlation coefficient (ρ) and p-values were determined using stats models in Python 3 or Matlab. Correlations between brain regions were visualized using network modeling in Matlab.

To investigate differences between the behaviors of mice in the open field we performed a compositional data analysis to account for the compositional nature of the data [88]. First, for each mouse, the set of fractions of time spent in each behavior were transformed into isometric log ratio coordinates using the R package ‘compositions’. In this coordinate space, differences between groups were analyzed using a nonparametric multivariate test (Wilks’ Lambda type test statistics) from the R package ‘npmv’. To identify the behaviors that differ between groups, we calculated the bootstrapped 95% confidence intervals (N=5000) of the log ratio differences between the cerebellar perturbed groups and mice without a cerebellar perturbation given CNO on day 1 for each behavior [88]. The same compositional data analysis was performed on the different control groups using the geometric mean values of all control groups combined as a reference group. The Matlab and R code used for the statistical analyses is published on Github (https://github.com/PrincetonUniversity/OF-ymaze-cfos-analysis).

## Data Availability

The dataset is available at Princeton data DOI: https://doi.org/10.34770/c9df-sc15 and https://brainmaps.princeton.edu/2022/01/verpeut-et-al-data-exploration-links/

All experimental and analysis code is available here: https://github.com/PrincetonUniversity/OF-ymaze-cfos-analysis

## Acknowledgements

We thank Henk-Jan Boele, Ben Deverett, Esteban Engel, Laura Lynch, Christina Matl, Dafina Pacuku, Tiffany Pham, Sanjeev Janarthanan, and Fred Uquillas for collaboration and discussion, Greg Horwitz for the L7 plasmid, and Archit Verma and Barbara Engelhardt for discussions on statistical testing. This work was supported by NIH R01 RS045193 and R01 MH115750 (S.W.) and the New Jersey Brain Injury Research Council (J.V.).

