## Supplemental figures for "Cerebellar contributions to a brainwide network for reversal learning"

**Corresponding authors

**This PDF file includes:**

Figures S1 to S14
Tables S1 to S3
Legends for Movies S1 to S3
Legends for Datasets S1

**Other supplementary materials for this manuscript include the following:**

Movies S1 to S3
Dataset S1

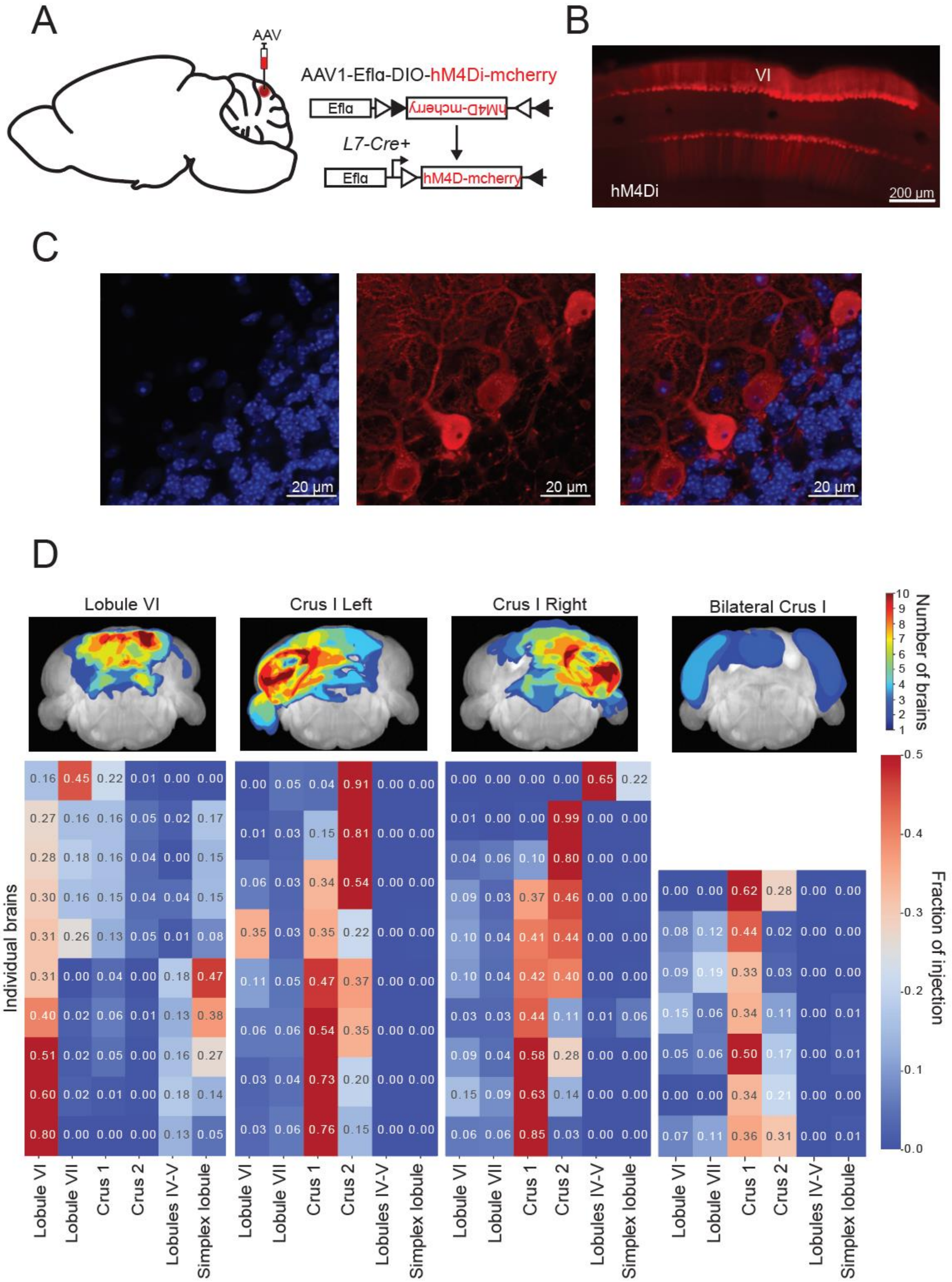


**S1 Fig. (Related to Fig 1) Selective expression of mCherry in Purkinje cells of the cerebellum.** (A) AAV carrying hM4Di fused to mCherry was injected into lobule VI or crus I of C57BL/6J mice along with AAV-L7-cre virus to allow for Purkinje cell-specific expression. (B) Sagittal section showing fluorescent expression (epifluorescent images at 40x) revealed by anti-RFP immunohistochemistry 3 weeks after injection in lobule VI. (C) Immunostaining confocal fluorescent images (63x) of lobule VI reveal hM4Di-mCherry expression in Purkinje cells. (D) Top: Whole-brain lightsheet averaged injection volumes calculated based on immunostaining of mCherry. Bottom: Calculated subset of individual brain volumes of the DREADD fraction injected per cerebellar lobule of interest.

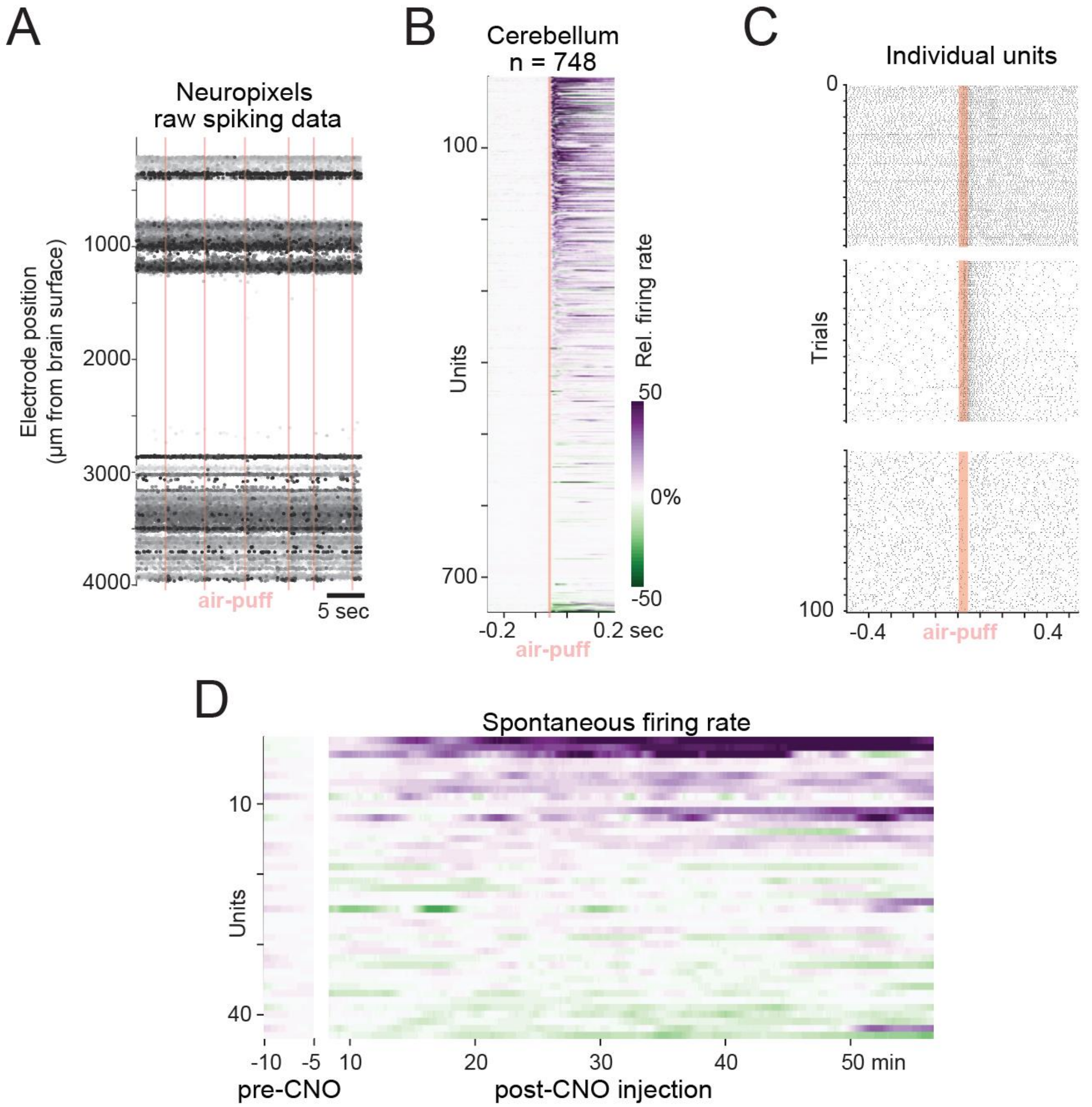


**S2 Fig. (Related to Fig 1) Neuropixels recordings of DREADD-expression in Purkinje cells.** (A) Raw spiking data on all 384 electrodes of the probe placed in cerebellum during a sensory stimulation with air puff. Stimulus time (40 ms, 1 pulse) is indicated with a red bar. (B) Normalized averaged responses of all sorted units to an air-puff stimulation. (C) Raster of spikes for 100 repetitions of the air-puff stimulation for three example units. Stimulus time (40 ms, 1 pulse) is indicated with a red bar. (D) Spontaneous firing rate of deep cerebellar nuclei neurons before and after CNO injection to mice with Purkinje cells expressing DREADD receptors.

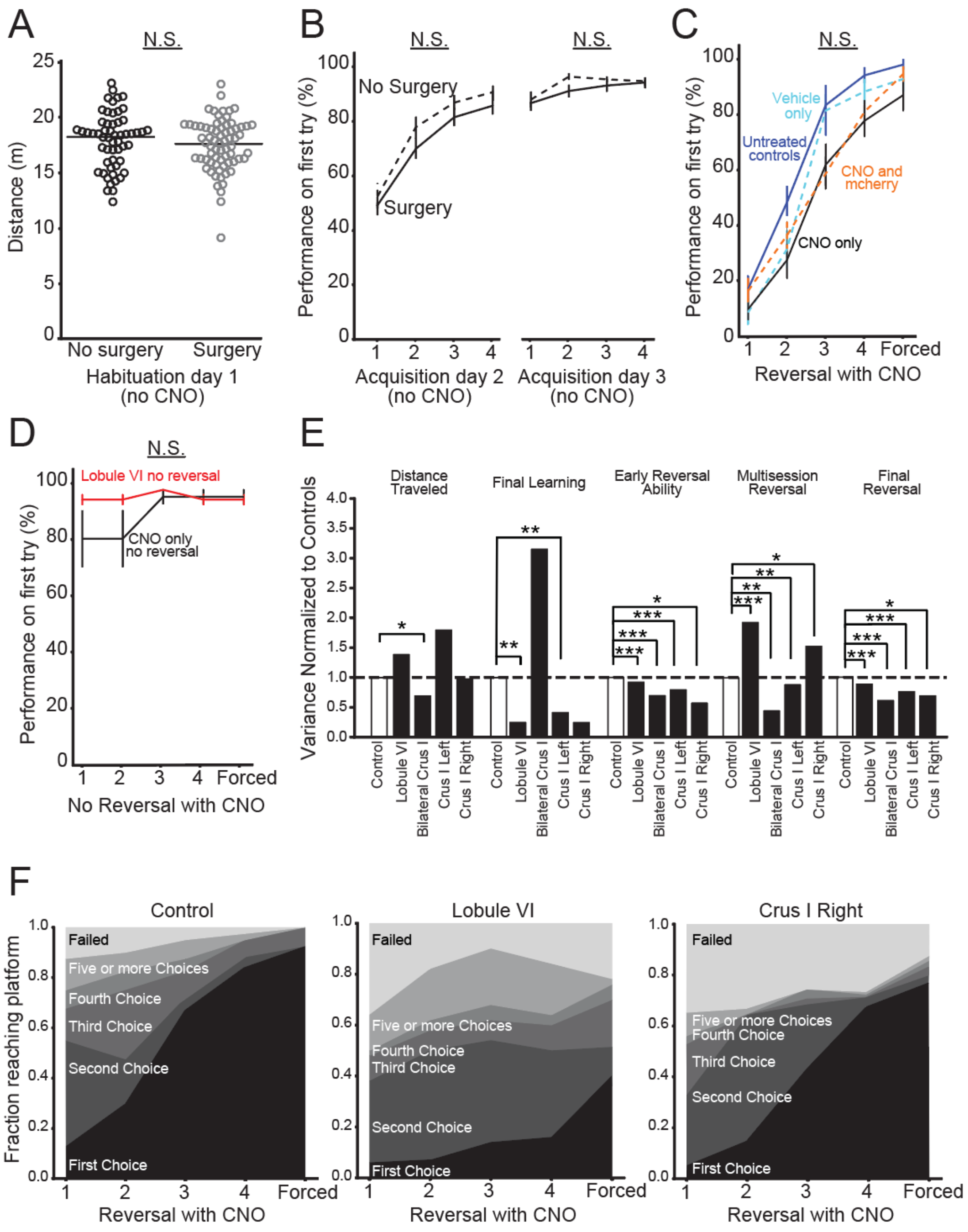


**S3 Fig. (Related to Fig 2)** Additional Y-maze metrics. (A) Surgery (DREADD experimental conditions and mCherry) had no effect on swimming total distance during habituation compared to no surgery (Untreated, Vehicle only, CNO only). (B) All animals were taught to swim to one single side of the Y-maze during acquisition sessions. Surgery had no effect on acquisition day 1 or 2. (C) To control for the unintended consequences of CNO, mice were given CNO or vehicle 20 minutes before testing. To control for potential AAV effects, animals received AAV with mCherry only and CNO during testing. No effects from AAV, CNO, or vehicle were found. (D) To test if activating the DREADD virus in lobule VI perturbed acquisition, a separate group of lobule VI mice were tested using a third acquisition day (no reversal) and compared to a CNO only no reversal group. No effects of CNO or DREADD perturbation were found in the no reversal condition. (E) Variance in distance traveled is less than during learning and reversal within experimental groups compared to untreated controls. (F) The number of choices required to find the platform in the Y-maze comparing experimental groups to CNO only control. Comparisons were made using one or two-way repeated measures ANOVA. Error bars indicate mean ± SEM. * $p < 0.05$, ** $p < 0.01$, *** $p < 0.001$

Control reversal over control no reversal

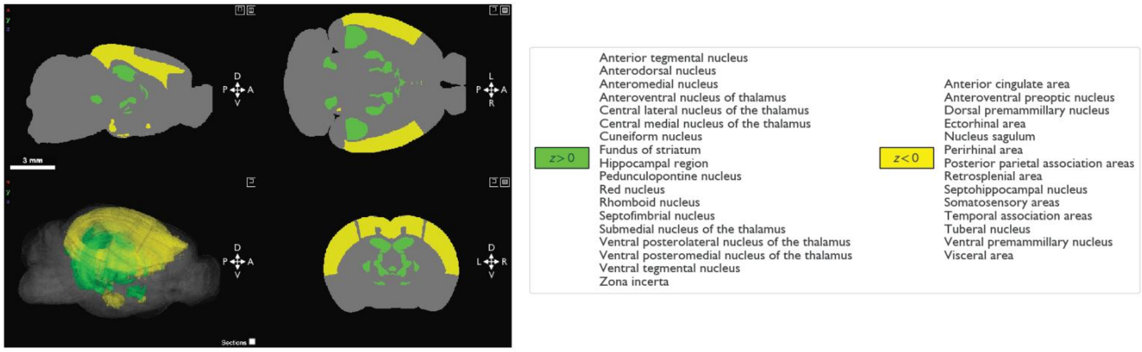


**S4 Fig. (Related to Fig 3) Reversal learning alters region-specific activity.** Whole-brain Neuroglancer visualization of significantly different regions in CNO only reversal compared to no reversal controls. Reversal resulted in different activated brain regions compared to acquisition. Green: increases, $p < 0.05$, Yellow: decreases, $p < 0.05$

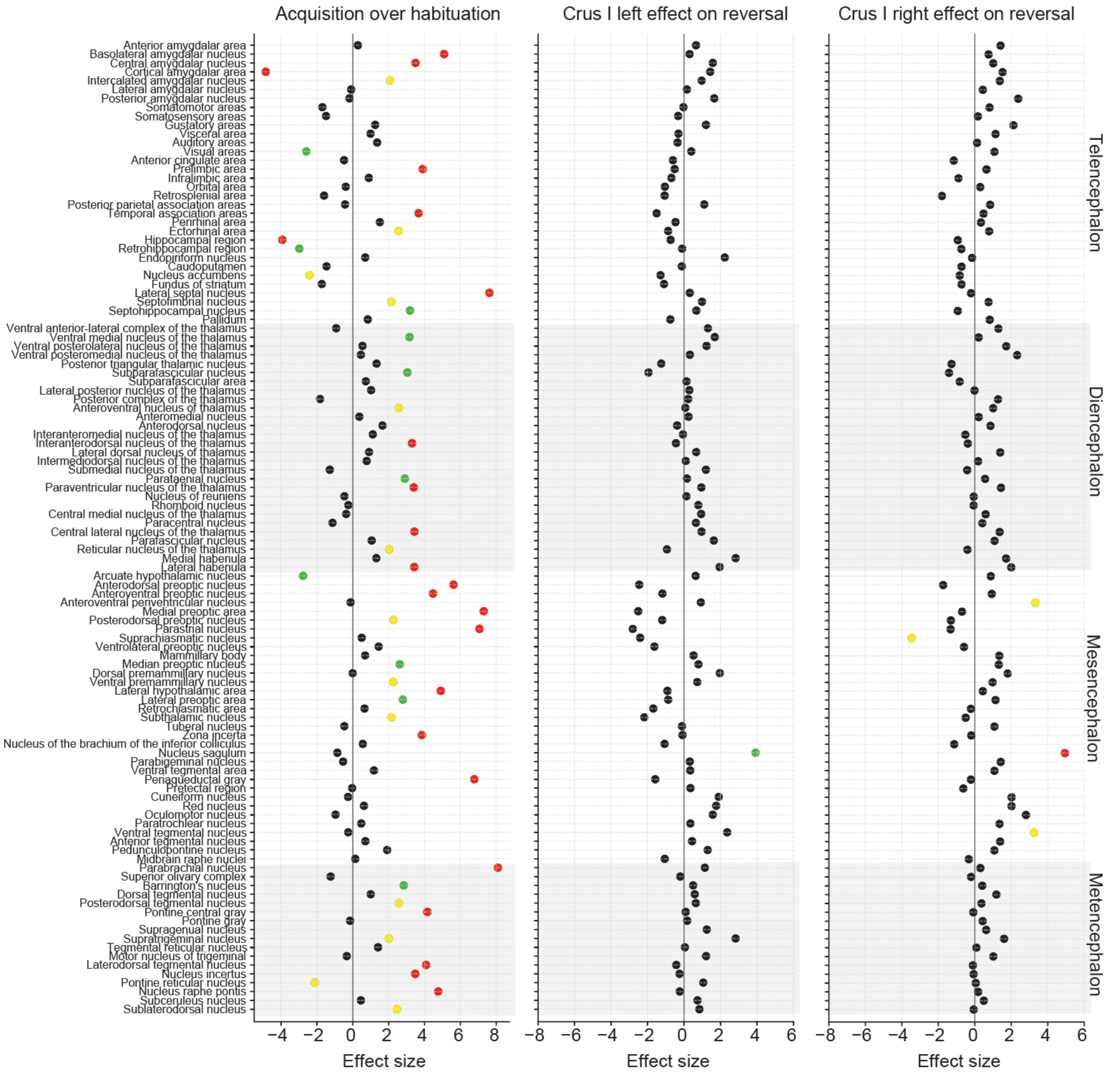


**S5 Fig. (Related to Fig 5) Additional comparisons from whole-brain imaging of c-Fos.** (Left) Comparisons between acquisition only and habituation only groups are plotted by effect size differences to find structures activated by the acquisition condition. Unilateral crus I left (middle) and crus I right (right) perturbation had little effect on c-Fos cell counts compared to CNO only control, which is consistent with a lack of a behavioral effect during reversal in the water Y-maze task. Comparisons were made using a negative binomial regression.Yellow: $p < 0.05$, Green: $p < 0.01$, Red: $p < 0.001$

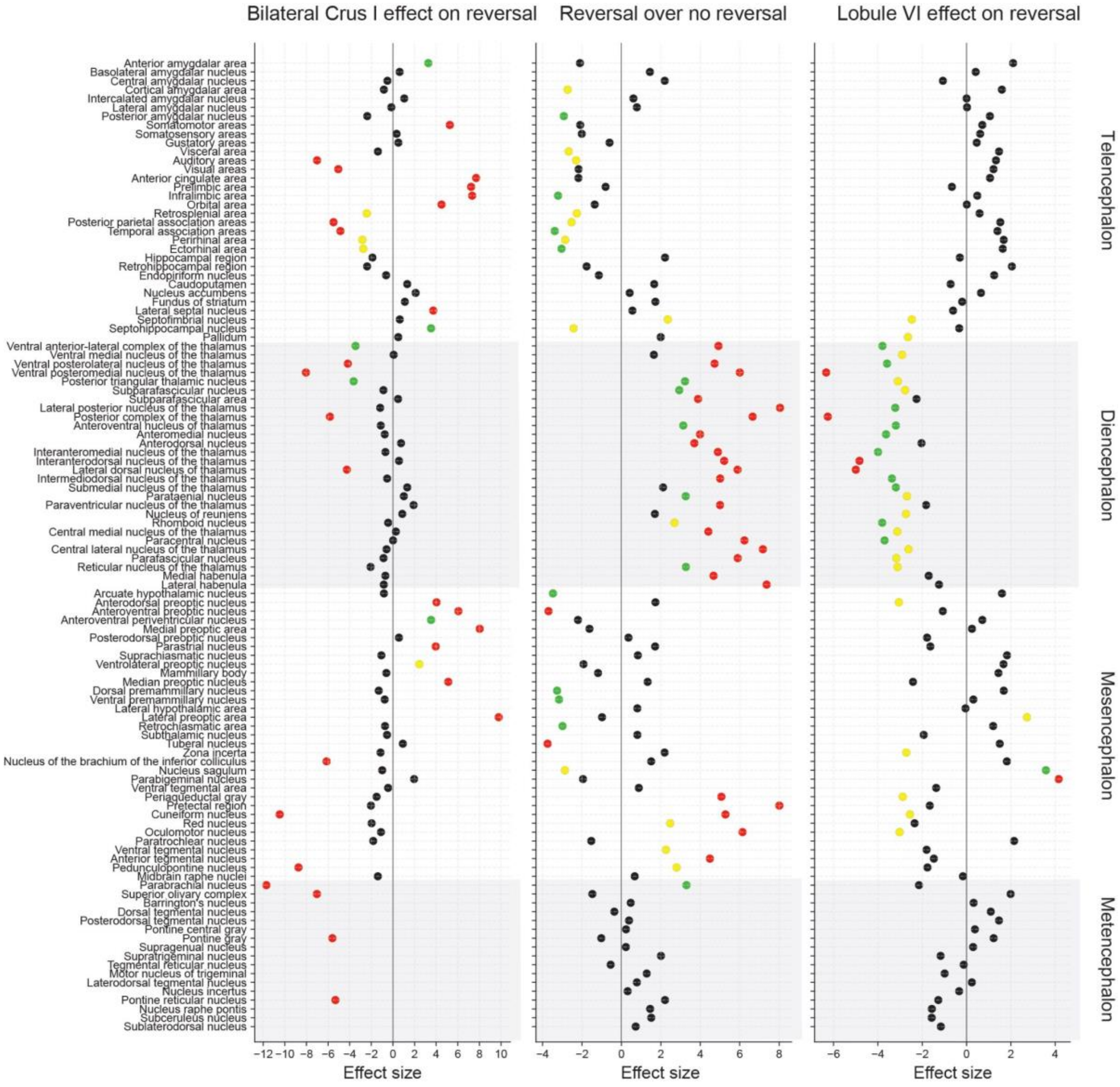


**S6 Fig. (Related to Fig 5) Brain-wide association study of activated c-Fos expression with distance swum as a variable.** Statistically-significant ($p < 0.05$) bilateral crus I (left), reversal (middle) lobule VI (right) structures compared to CNO only controls. By adding distance swum as a variable in our analysis we find most structures remain significant suggesting that swimming is a separate neural network to Y-maze reversal. Yellow: $p < 0.05$, Green: $p < 0.01$, Red: $p < 0.001$

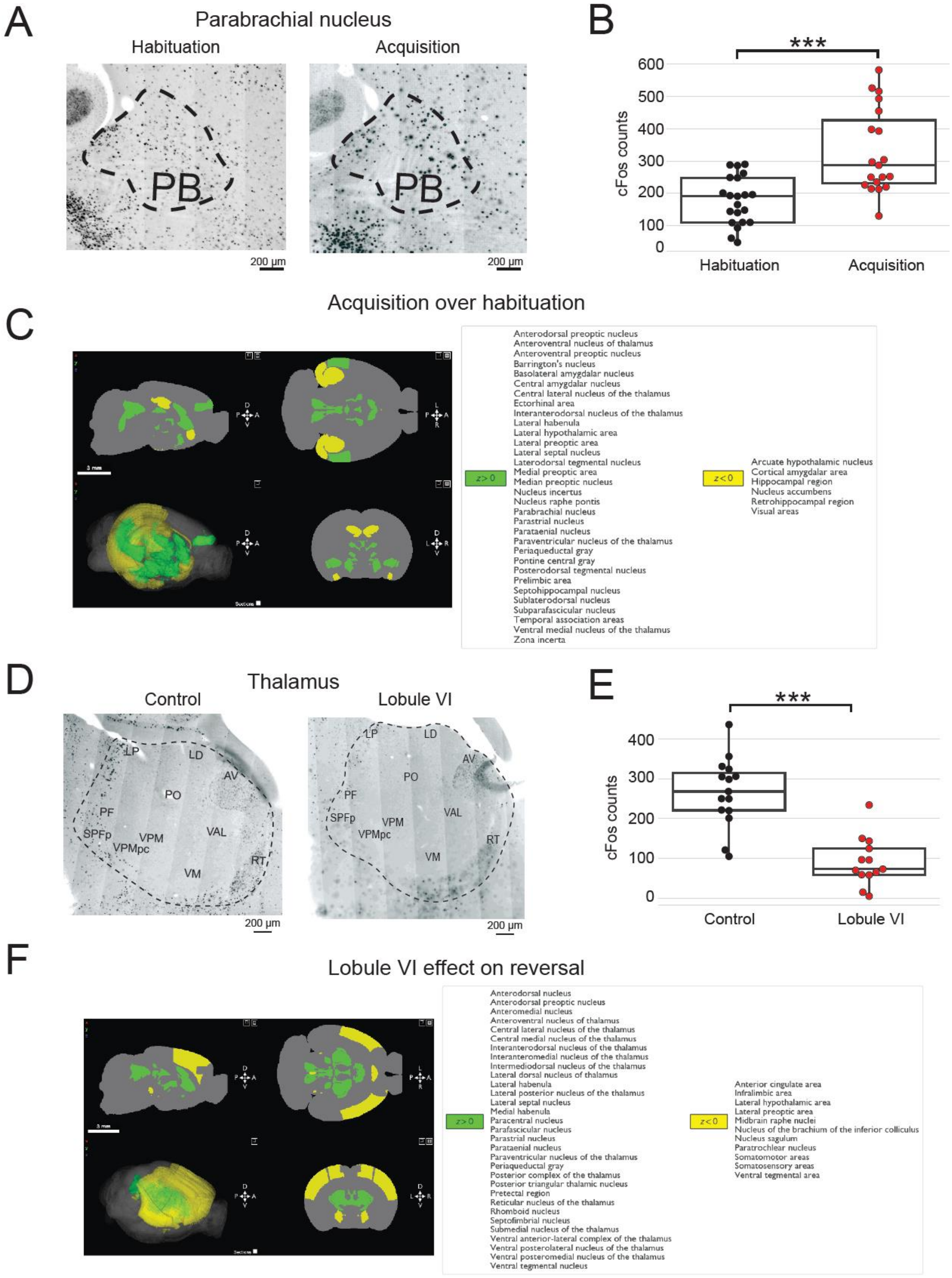
A
Parabrachial nucleus
Habituation
Acquisition
PB
PB
200 µm
200 µm
B
***
cFos counts
600
500
400
300
200
100
0
Habituation
Acquisition
Acquisition over habituation
C
z > 0
Anterodorsal preoptic nucleus
Anteroventral nucleus of thalamus
Anteroventral preoptic nucleus
Barrington's nucleus
Basolateral amygdalar nucleus
Central amygdalar nucleus
Central lateral nucleus of the thalamus
Ectorhinal area
Interanterodorsal nucleus of the thalamus
Lateral habenula
Lateral hypothalamic area
Lateral preoptic area
Lateral septal nucleus
Laterodorsal tegmental nucleus
Medial preoptic area
Median preoptic nucleus
Nucleus incertus
Nucleus raphe pontis
Parabrachial nucleus
Parastrial nucleus
Parataenial nucleus
Paraventricular nucleus of the thalamus
Periaqueductal gray
Pontine central gray
Posterodorsal tegmental nucleus
Prelimbic area
Septohippocampal nucleus
Sublaterodorsal nucleus
Subparafascicular nucleus
Temporal association areas
Ventral medial nucleus of the thalamus
Zona incerta
z < 0
Arcuate hypothalamic nucleus
Cortical amygdalar area
Hippocampal region
Nucleus accumbens
Retrohippocampal region
Visual areas
D
Thalamus
Control
Lobule VI
LP
LD
AV
PO
PF
SPFp
VPM
VPMpc
VAL
VM
RT
200 µm
200 µm
E
***
cFos counts
400
300
200
100
0
Control
Lobule VI
F
Lobule VI effect on reversal
z > 0
Anterodorsal nucleus
Anterodorsal preoptic nucleus
Anteromedial nucleus
Anteroventral nucleus of thalamus
Central lateral nucleus of the thalamus
Central medial nucleus of the thalamus
Interanterodorsal nucleus of the thalamus
Interanteromedial nucleus of the thalamus
Intermediodorsal nucleus of the thalamus
Lateral dorsal nucleus of thalamus
Lateral habenula
Lateral posterior nucleus of the thalamus
Lateral septal nucleus
Medial habenula
Paracentral nucleus
Parafascicular nucleus
Parastrial nucleus
Parataenial nucleus
Paraventricular nucleus of the thalamus
Periaqueductal gray
Posterior complex of the thalamus
Posterior triangular thalamic nucleus
Pretectal region
Reticular nucleus of the thalamus
Rhomboid nucleus
Septofimbrial nucleus
Submedial nucleus of the thalamus
Ventral anterior-lateral complex of the thalamus
Ventral posterolateral nucleus of the thalamus
Ventral posteromedial nucleus of the thalamus
Ventral tegmental nucleus
z < 0
Anterior cingulate area
Infralimbic area
Lateral hypothalamic area
Lateral preoptic area
Midbrain raphe nuclei
Nucleus of the brachium of the inferior colliculus
Nucleus sagulum
Paratrochlear nucleus
Somatomotor areas
Somatosensory areas
Ventral tegmental area

**S7 Fig. (Related to Fig 3-5) Acquisition and lobule VI perturbation activate an unique set of brain regions in the water Y-maze task.** (A) Conventional immunohistochemical staining of c-Fos in the parabrachial nucleus (PB) in habituation only versus acquisition only groups. (B) Acquisition only mice had increased c-Fos staining in PB. Scale bar 200 μm (C) Whole-brain Neuroglancer visualization of significantly different regions in acquisition circuits. (D) Conventional immunohistochemical staining of c-Fos in the thalamus of Lobule VI compared to CNO only. Lateral posterior nucleus (LP), lateral dorsal nucleus (LD), anteroventral nucleus (AV), posterior complex (PO), ventral anterior-lateral complex (VAL), ventral medial nucleus (VM), reticular nucleus (RT), ventral posteromedial nucleus (VPM), ventral posteromedial nucleus parvicellular part (VPMpc), subparafascicular parvicellular part (SPFp), parafascicular nucleus (PF). (E) Lobule VI mice had significantly less c-Fos in thalamic regions compared to CNO only. (F) Whole-brain Neuroglancer visualization of significantly different regions in lobule VI circuits. Green: increases, $p < 0.05$, Yellow: decreases, $p < 0.05$, *** $p < 0.001$

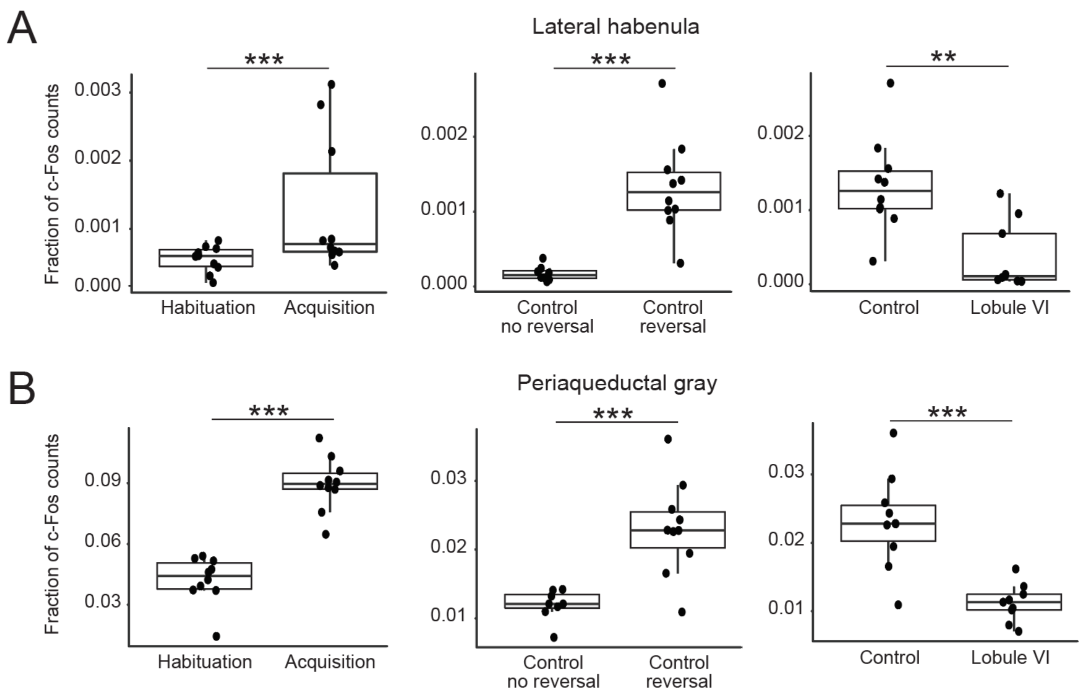


**S8 Fig. (Related to Fig 3-5) c-Fos neural activation in lateral habenula and periaqueductal gray.** (A) Y-maze acquisition (left) and reversal learning in CNO only controls (middle) significantly increased c-Fos in the lateral habenula. Lobule VI perturbation reduced c-Fos counts in the lateral habenula compared to reversal learning (right). (B) In the periaqueductal gray, Y-maze acquisition (left) and reversal learning in CNO only controls (middle) significantly increased c-Fos counts. Lobule VI perturbation reduced c-Fos counts in the periaqueductal gray compared to reversal learning (right). Comparisons were made using a pairwise t-test. ** $p < 0.01$, *** $p < 0.001$

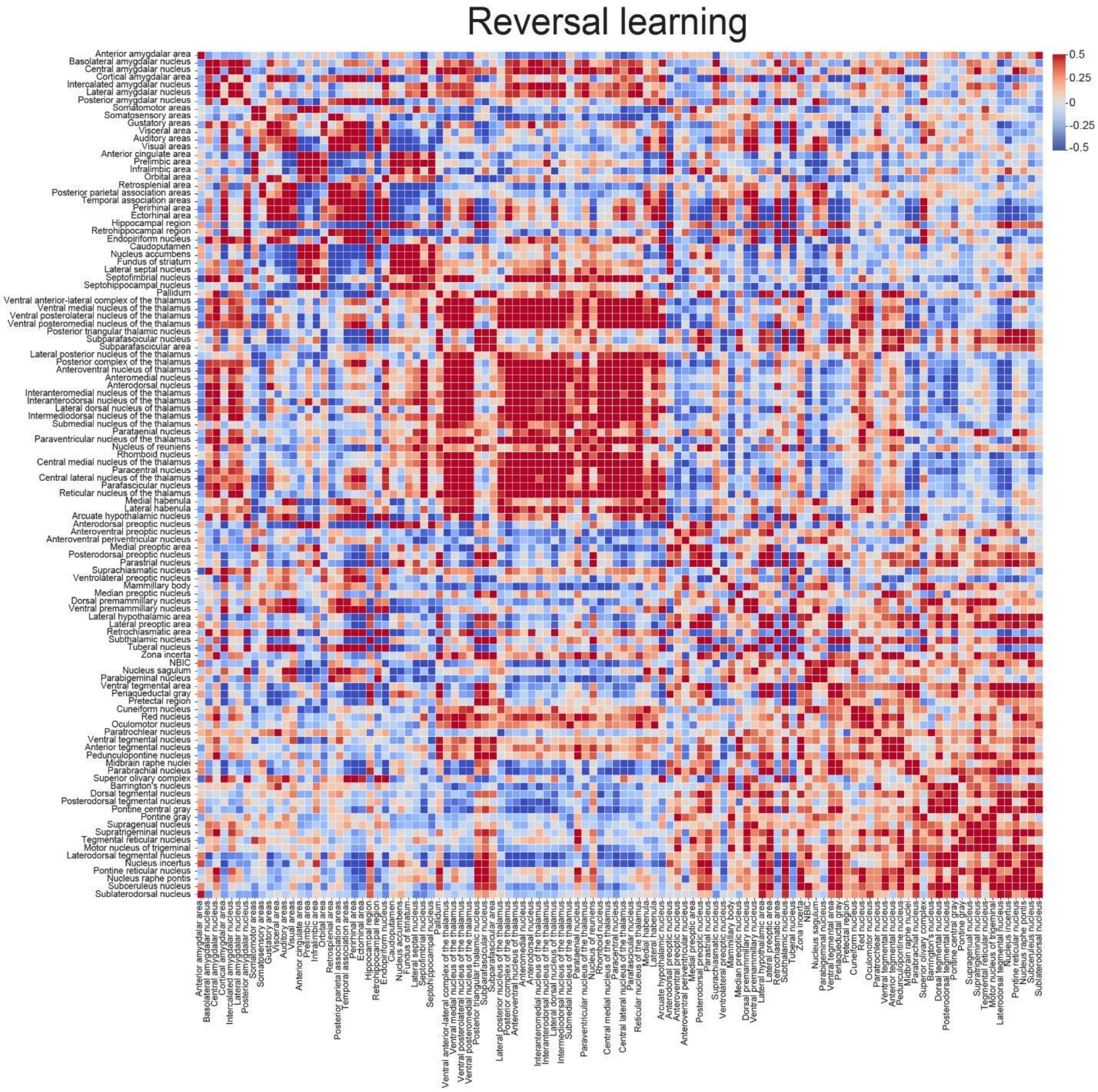


**S9 Fig. (Related to Fig 6) Reversal learning correlation matrix.** Inter-region connections for c-Fos expression in reversal learning. Strength of correlation reflected in scale bar (Spearman's p).

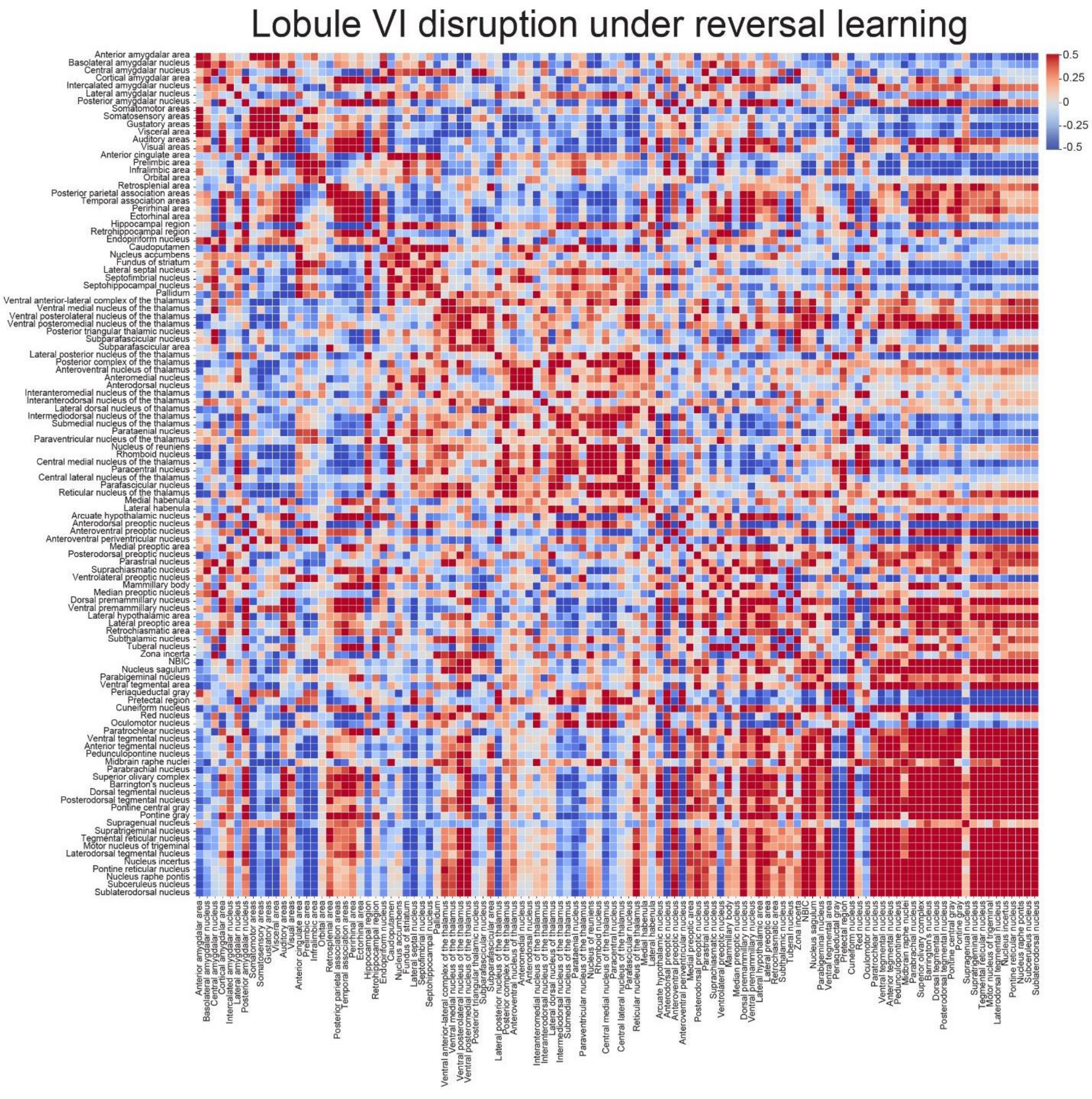


**S10 Fig. (Related to Fig 6). Lobule VI disruption in Y-maze reversal.** Inter-region connections for c-Fos expression in lobule VI disruption. Strength of correlation reflected in scale bar (Spearman’s ρ).

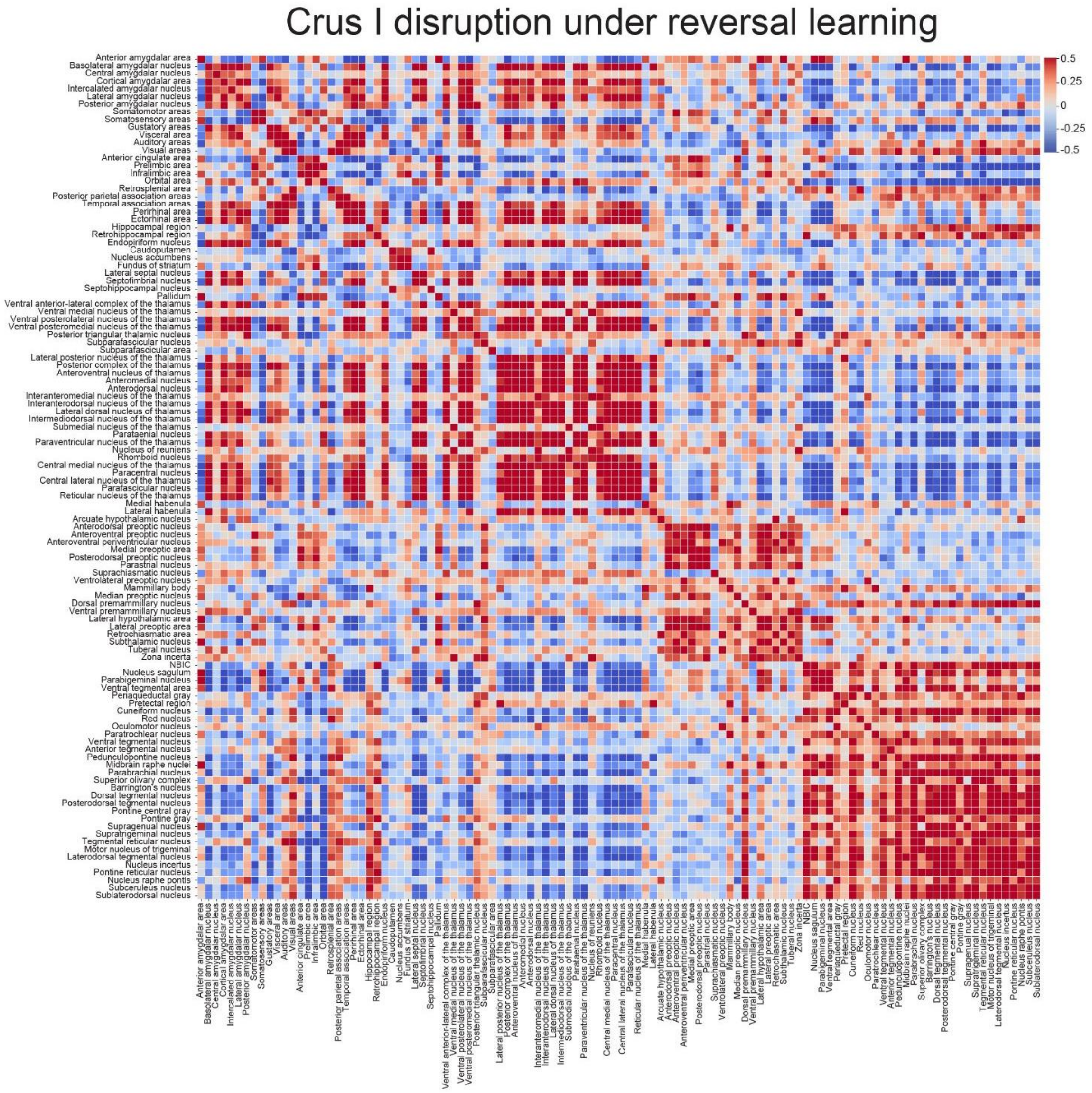


**S11 Fig. (Related to Fig 6). Crus I disruption in Y-maze reversal.** Inter-region connections for c-Fos expression in crus I disruption. Strength of correlation reflected in scale bar (Spearman's p).

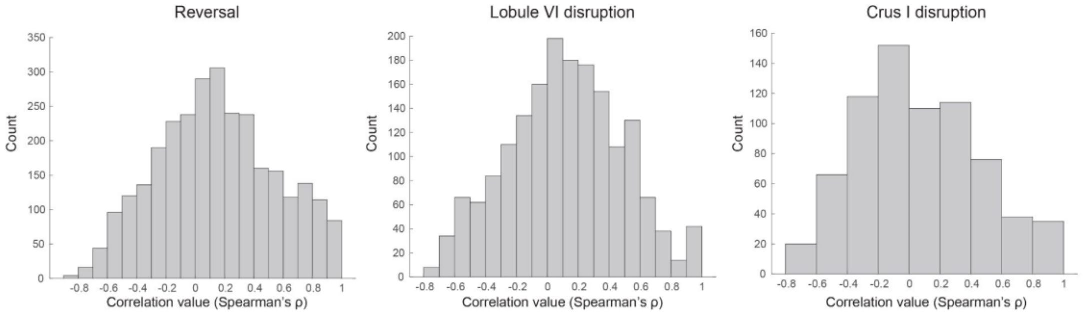


**S12 Fig. (Related to Fig 6) Histogram analysis of inter-region correlations.** Correlation (Spearman's ρ) histograms for significant regions found in brainwide comparisons of groups at $p < 0.05$ for reversal (left), lobule VI disruption (middle), and combined crus I disruption (right).

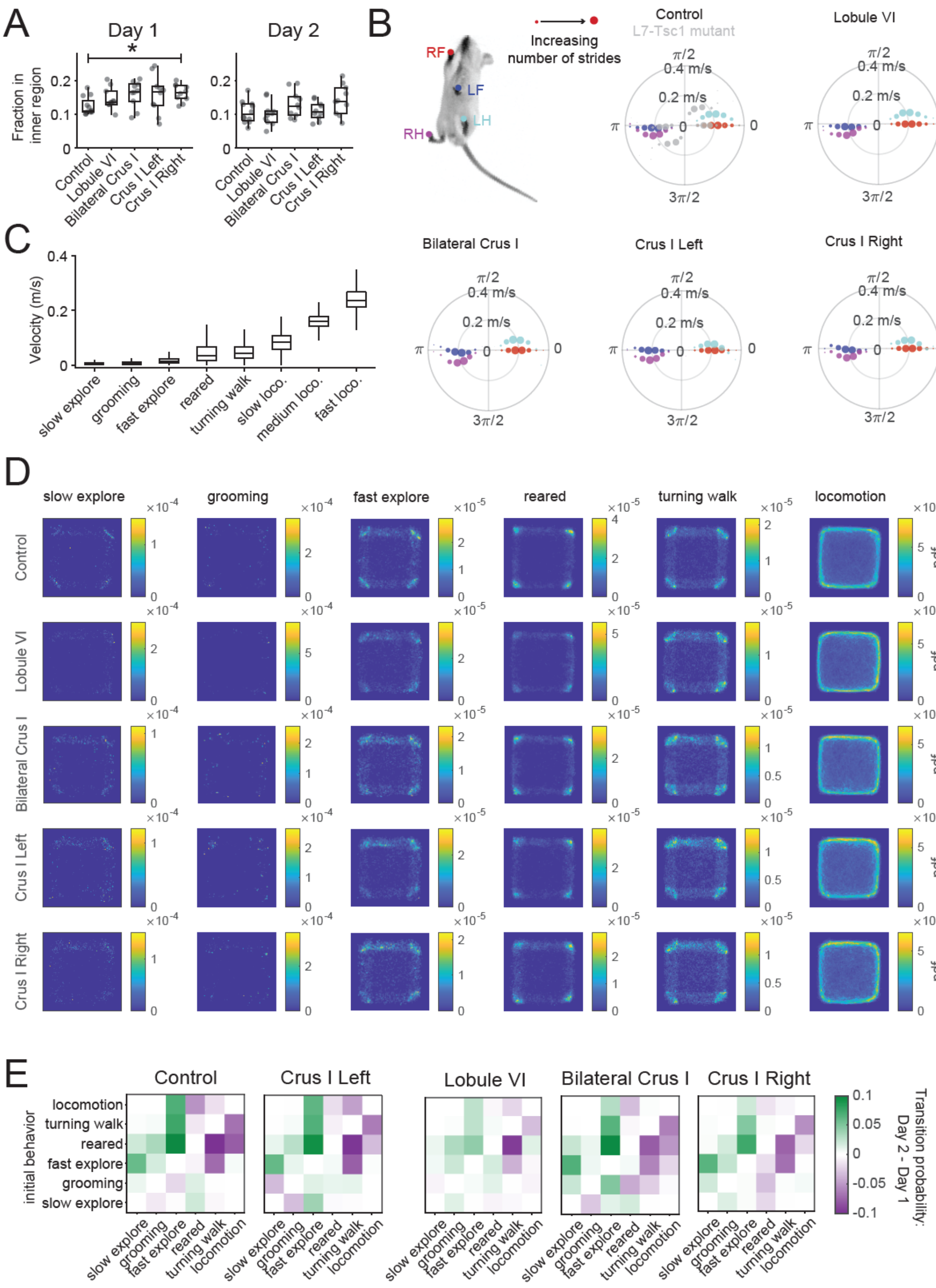
A
Day 1
Day 2
Fraction in inner region
Control
Lobule VI
Bilateral Crus I
Crus I Left
Crus I Right
B
RF
LF
LH
RH
Increasing number of strides
Control
L7-Tsc1 mutant
Lobule VI
Bilateral Crus I
Crus I Left
Crus I Right
0.4 m/s
0.2 m/s
C
Velocity (m/s)
slow explore
grooming
fast explore
reared
turning walk
slow loco.
medium loco.
fast loco.
D
slow explore
grooming
fast explore
reared
turning walk
locomotion
pdf
E
Control
Crus I Left
Lobule VI
Bilateral Crus I
Crus I Right
initial behavior
locomotion
turning walk
reared
fast explore
grooming
slow explore
Transition probability: Day 2 - Day 1

**S13 Fig. (Related to Fig 7) Lobule-specific alterations of spontaneous behavior in open field.** (A) Fraction of time spent in the inner region of the arena. Comparisons of the groups within each day to CNO only control were made using a Kruskal-Wallis test, followed up by pairwise comparisons using Wilcoxon rank sum exact tests with Benjamini-Hochberg correction. (B) Limb coordination during locomotion is not impaired by cerebellar perturbations. The maximal values of the paw positions in an animal-centered coordinate system were assumed at similar phases during single strides of locomotion. For comparison altered limb coordination in L7-Tsc1 mutant mice is shown from previously published experiments (7). (C) Centroid velocities during different behaviors. (D) The spatial distributions of occurrences of the different behaviors in the open field arena are similar for each experimental group compared to CNO only control. Data from the recording on day 1 was considered. (E) Lobule VI-perturbed CNO only control mice showed the smallest differences in the state transition probabilities across days. * $p < 0.05$

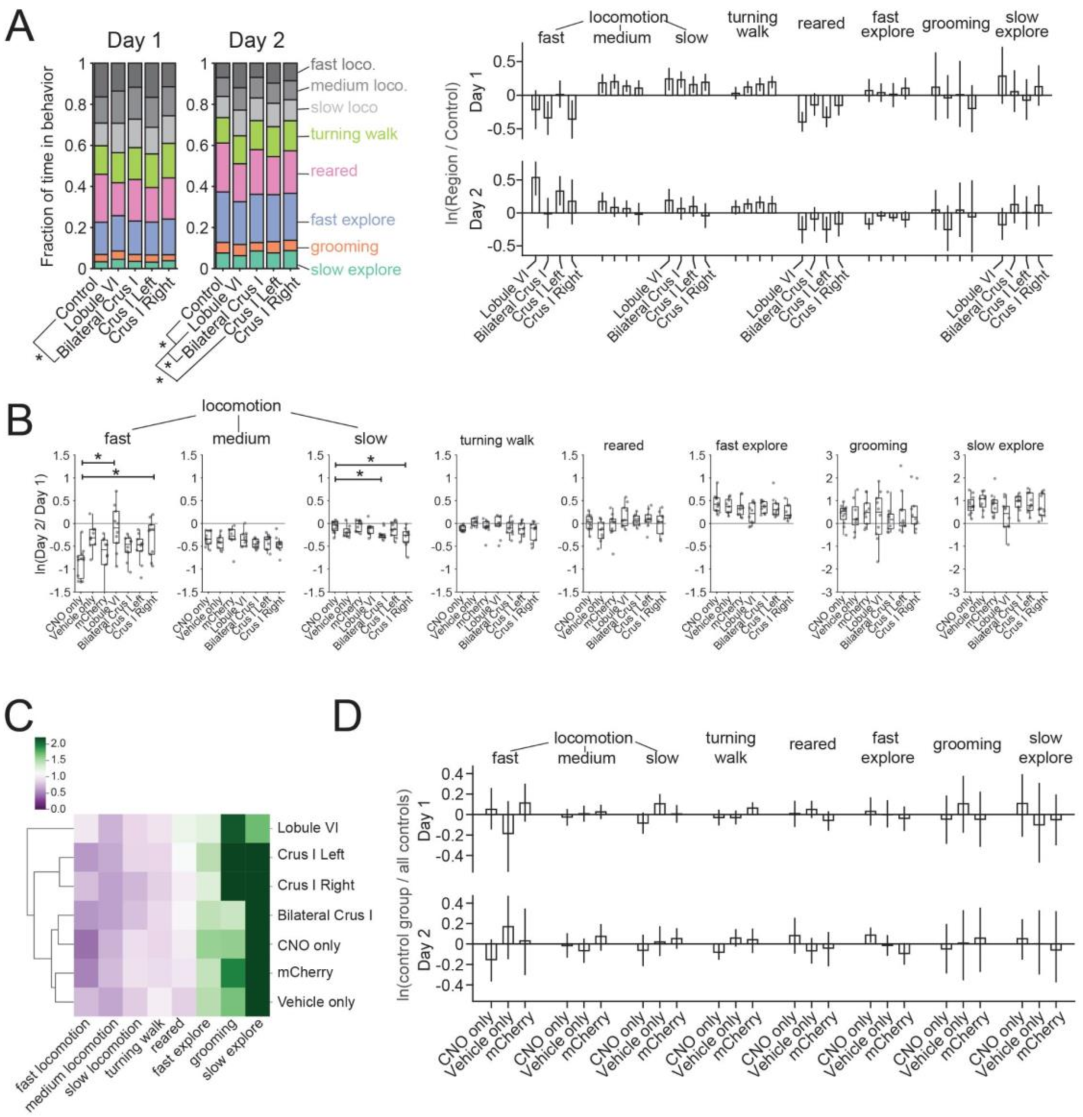
A
Day 1
Day 2
Fraction of time in behavior
fast loco.
medium loco.
slow loco
turning walk
reared
fast explore
grooming
slow explore
Control
Lobule VI
Bilateral Crus I
Crus I Left
Crus I Right
locomotion
fast
medium
slow
turning walk
reared
fast explore
grooming
slow explore
ln(Region / Control)
B
ln(Day 2/ Day 1)
CNO only
Vehicle only
mCherry
C
fast locomotion
medium locomotion
slow locomotion
turning walk
reared
fast explore
grooming
slow explore
D
ln(control group / all controls)

**S14 Fig. (Related to Fig 7) Comparison of spontaneous behavior in control groups.** (A) The fractions of time spent in each of the eight behaviors (compositional means) differ significantly between lobule VI-perturbed mice and the CNO only control group on day 2 (nonparametric multivariate test on ilr-transformed fractions, ssnonpartest from R package npmv, Wilks' Lambda type statistic). This difference can be mainly accounted for by the large amount of time spent in fast locomotion on the second day compared to the control, quantified by the log ratio differences between the cerebellar perturbed groups and the control (error bars indicate bootstrapped 95% confidence intervals, N = 5000, percentile bootstrap) (B) Cerebellar perturbations of lobule VI, crus I right and bilateral crus I had an effect on the adaptation of the occupancy in slow and fast locomotion states across days compared to CNO only control (Kruskal-Wallis tests for each behavior, pairwise comparisons using Wilcoxon rank sum exact test, Benjamini-Hochberg correction). (C) The state occupancies (Day 2/Day 1) for all experimental and control groups show clustering between controls compared to experimental groups. Crus I right and left cluster together and have similar state occupancies. Lobule VI-perturbed mice are the most different from all other groups, emphasizing that the lack of adaptation creates an unique phenotype. (D) All control groups show a similar change of behavioral occupancies across days (Kruskal-Wallis tests for each behavior, pairwise comparisons using Wilcoxon rank sum exact test, Benjamini-Hochberg correction). * $p < 0.05$

**S1 Table:** Y-maze distance swum metrics (m).

| | Habituation only | Acquisition only | Control reversal | Lobule VI | Crus I bilateral | Crus I right | Crus I left | CNO no reversal | Lobule VI no reversal |
|---|---|---|---|---|---|---|---|---|---|
| N | 10 | 10 | 22 | 16 | 7 | 25 | 26 | 7 | 10 |
| Mean | 54.6 | 23.2 | 20.2 | 22.7 | 25.8 | 22.3 | 22.3 | 17.1 | 16.9 |
| SD | 12.2 | 6.2 | 6.0 | 8.6 | 7.4 | 6.8 | 7.4 | 4.8 | 3.6 |

**S2 Table:** Key Resources Table

| Reagent type (species) or resource | Designation | Source or reference | Identifiers |
|---|---|---|---|
| strain, strain background | Mouse :C57BL/6J | The Jackson Laboratory, Bar Harbor, ME | Stock#: 00664 \|Black6 https://www.jax.org/strain/000664 |
| strain, strain background | Mouse: *Pcp2*$^{Cr}$ | The Jackson Laboratory, Bar Harbor, ME | Stock#: 004146 https://www.jax.org/strain/004146 |
| antibody | anti-GFP chicken | Aves Labs | Cat#GFP-1020; RRID: AB_10000240 |
| antibody | goat anti-chicken IgY (H+L) secondary antibody, Alexa Fluor 647 | ThermoFisher | Cat#A-21449; RRID: AB_2535866 |
| recombinant DNA reagent | AAV1-Eflα-DIO-hM4D(Gi)-mCherry-WPRE-hGHpA;AAV8-Eflα-DIO-mCherry-WPRE-hGHpA | Princeton Vector Core | |
| recombinant DNA reagent | AAV1-sL7-Cre-HA-WPRE-hGH-pA | Princeton Vector Core | |
| chemical compound, drug | Clozapine-N-oxide (CNO) | NIMH Chemical Synthesis and Drug Supply Program | Cat# 34233-69-7 |
| chemical compound | ProLong™ Diamond Antifade Mountant | ThermoFisher | Cat# P36961 |
| chemical compound, drug | 15% D-mannitol | SIGMA-ALDRICH | Cat# M4125 |
| chemical compound | DPBS | ThermoFisher | Cat#14190136 |
| chemical compound, drug | white tempera paint | Artmind, Tempera Paint | Cat#10091773 |
| chemical compound, drug | Rimadyl [carprofen] | Zoetis, Florham Park, NJ | http://www.zoetisus.com |
| chemical compound | cholera toxin subunit B (CT-B), AlexaFluor 488 Conjugate | ThermoFisher | Cat# C34775 |
| chemical compound, drug | ketamine/xylazine | Met-Vet International/Akorn | RXV CIII (3N)/Cat# 59399-111-50 |
| chemical compound, drug | Euthasol | Met-Vet International/Virbac | RXV CIII (3N)/ RXEUTHASOL |
| chemical compound, | CM-DiI | Thermofisher | Cat # C7000 |

| | | | |
|---|---|---|---|
| software, algorithm | Illustrator CS | Adobe | |
| | Excel | Microsoft | |
| | MATLAB R2020b | MathWorks | |
| | Python 2.7.14/3.8.6 | Python | |
| | R 4.0.3 | R | |
| | ImageJ | NIH | |
| | Neuroglancer | (Google WebGL-based viewer) | |
| | Allen Brain Atlas | (Oh et al., 2014). | http://www.brain-map.org |
| Resource Availability | Data for this manuscript | Princeton database | https://doi.org/10.34770/c9df-sc15 |
| | Data from Klibaite et al., 2022 | Princeton database | https://dataspace.princeton.edu/handle/88435/dsp012j62s793m |
| | Code | Github | https://github.com/PrincetonUniversity/OF-ymaze-cfos-analysis |

**S3 Table:** Control and experimental conditions per figure.

| Control group | Experimental group | Figure |
|---|---|---|
| | Lobule VI | Fig 1C-G, Fig S1B-C, Fig S2A-D, Fig S10 |
| CNO only | Lobule VI | Fig 4C, Fig S3D-F, Fig 7A, Fig 7C, Fig 7E, Fig S7D-F |
| CNO only | Lobule VI, Bilateral Crus I, Crus I Left, Crus I Right | Fig 2B, Fig 6, Fig S3A-B and D-E, Fig 7D, S9 Fig, S12 Fig, S13A Fig, S13D-E Fig, S14A Fig |
| CNO only | Lobule VI, Bilateral Crus I | Fig 2C-D, Fig 5A |
| CNO only | Crus I Left, Crus I Right | Fig S5 |
| CNO only, CNO only no reversal | | Fig 3C-D, Fig S4, Fig S8, Fig S9 |
| Lobule VI no reversal | Lobule VI reversal | |
| CNO only, CNO only no reversal | Lobule VI, Bilateral Crus I | Fig 5B-C, Fig S6 |
| | Lobule VI, Crus I Left, Crus I Right | Fig S1D |
| CNO only, Vehicle only, mCherry | | Fig S4C and D (right), S14D Fig |
| CNO only, Vehicle only, mCherry | Lobule VI, Bilateral Crus I, Crus I Left, Crus I Right | S14B-C Fig |
| CNO only, Vehicle only, mCherry, Untreated only | | Fig S3C |
| Acquisition, Habituation | | Fig S5, Fig S7A-C |
| Acquisition, Habituation, CNO only, CNO no reversal | Lobule VI | Fig S8 |
| | Bilateral Crus I, Crus I Left, Crus I Right | Fig S11 |

**Movie S1 (separate file).**

1) S1 Movie. Y-maze task. Example video recording of mouse performing Y-maze acquisition, reversal with CNO only, and lobule VI perturbed mouse attempting reversal.
2) S2 Movie. Whole brain lightsheet. Example movie of a cleared brain using iDISCO+, c-Fos staining, clearmap cell counts, and neuroglancer overlay of multiple thalamic regions.
3) S3 Movie. Semi-supervised behavioral classification. Example video recording of mouse in open field (left), zoomed-in version with LEAP labels (middle) and corresponding ethogram (right).

**Dataset S1 (separate file).**

1) S1 Data. Effects of experimental perturbations on c-Fos cell counts. Contrasts include acquisition versus habituation, reversal learning versus acquisition, and lobule VI perturbation via DREADDs against CNO only. For each brain region and contrast, negative binomial (NB) regression was performed with natural log of total counts across all regions as an offset. Estimate: NB slope coefficient for effect of treatment (log-ratio scale), Fold change: exponentiated coefficient (ratio scale), Std. Error: standard error of estimate, z value: estimate divided by standard error (also known as effect size), Pr(>|z|): raw p-value for effect, fdr_adj_pval: p-value adjusted for false discovery rate, status: whether NB fitting routine encountered numerical problems.
